# Encapsulation of Select Violacein Pathway Enzymes in the 1,2-Propanediol Utilization Bacterial Microcompartment to Divert Pathway Flux

**DOI:** 10.1101/2024.11.15.623819

**Authors:** Brett Jeffrey Palmero, Emily Gamero, Niall M. Mangan, Danielle Tullman-Ercek

## Abstract

A continual goal in metabolic engineering is directing pathway flux to desired products and avoiding loss of pathway intermediates to competing pathways. Encapsulation of the pathway is a possible solution, as it creates a diffusion barrier between pathway intermediates and competing enzymes. It is hypothesized that bacteria use organelles known as bacterial microcompartments - proteinaceous shells encapsulating a metabolic pathway - for this purpose. We aim to determine to what degree this hypothesized benefit is conferred to encapsulated pathways. To this end, we used bacterial microcompartments to encapsulate select enzymes from the violacein pathway, which is composed of five enzymes that produce violacein as the main product and deoxyviolacein as a side product. Importantly, we studied the pathway in a cell-free context, allowing us to hold constant the concentration of unencapsulated and encapsulated enzymes and increase our control over reaction conditions. The VioE enzyme is a branch point in that it makes the precursor for both violacein and deoxyviolacein, the VioC enzyme is required for production of deoxyviolacein, and the VioD enzyme is required for violacein production. When we encapsulated VioE and VioC and left VioD unencapsulated, the product profile shifted toward deoxyviolacein and away from violacein compared to when VioC and VioD were both unencapsulated. This work provides the first fully quantitative evidence that microcompartment-based encapsulation can be used to divert pathway flux to the encapsulated pathway. It provides insight into why certain pathways are encapsulated natively and could be leveraged for metabolic engineering applications.

## Introduction

Metabolic engineering has the potential to utilize biological systems to make industrially relevant products from cheap, renewable feedstocks with higher energy efficiency (1–4). Multiple groups have successfully utilized metabolic pathways to produce commodity chemicals, but few have reached commercialization, often due to low yields (5–7). A variety of factors drive this poor output, including loss of pathway flux to off-target reactions and cofactor imbalances for the pathway of interest. These challenges can be further exacerbated by toxic intermediate accumulation within the cell, reducing cellular fitness.

Several of these challenges are not unique to metabolic engineering; bacteria natively experience similar challenges including reduced cell viability due to toxic intermediate build-up and pathway cofactor imbalances. Many bacterial phyla harbor a specialized organelle known as a bacterial microcompartment (MCP), which is hypothesized to overcome these challenges (8–12). MCPs consist of an outer porous protein shell and an enzymatic core of encapsulated metabolic enzymes. They can be classified as carboxysomes or metabolosomes depending on whether they encapsulate anabolic or catabolic reactions, respectively (11). Encapsulation of native metabolic pathways within metabolosomes has been shown to retain toxic pathway intermediates and provide a private cofactor pool for the encapsulated pathway (13–18). Both benefits rely on the presence of a diffusion barrier around the pathway, which could be used in metabolic engineering to sequester intermediates of a pathway of interest away from competing enzymatic pathways, reducing off-target reactions. One of the best-characterized metabolosomes is the 1,2-propanediol utilization MCP (Pdu MCP) which breaks down 1,2-propanediol (1,2-PD) into propionate for bacteria to use as a carbon and energy source. The native Pdu pathway produces propionaldehyde, a toxic intermediate that can build up and inhibit cellular fitness if the pathway enzymes are not encapsulated in the Pdu MCP (19–25). Experiments show that cellular fitness is improved by sequestering the toxic intermediate propionaldehyde in the Pdu MCP, a finding that is supported with kinetic models (19, 21). It has been long hypothesized that intermediate sequestration in MCPs can also be used to divert pathway intermediates to the encapsulated enzymes and avoid competing reactions, but to date, there are no studies that have quantitatively examined this hypothesis (26, 27).

Multiple groups have investigated how encapsulating heterologous multi-enzyme pathways in MCPs and structures based on MCP shell components affects overall metabolic pathway performance. Lawrence *et al*. were the first to encapsulate a heterologous pathway in an MCP and achieved a 1.63-fold increase in ethanol production by encapsulating pyruvate decarboxylase and alcohol dehydrogenase (28). In another study, Lee *et al.* encapsulated four enzymes involved in 1,2- PD production in an MCP and saw an approximately 3-fold increase in 1,2-PD yield compared to when using unencapsulated enzymes (29). Surprisingly, they observed that the highest yield of 1,2- PD came from enzymes that were colocalized in a shell-free aggregate but not encapsulated in Pdu MCP shells. Additionally, Kirst *et al*. used the SpyCatcher/SpyTag system to encapsulate a formate utilization pathway in a modified *Haliangium ochraceum* MCP (30). This pathway consisted of phosphotransacetylase and pyruvate formate lyase and was capable of recycling acetyl-CoA into CoA. Both enzymes were active when encapsulated, suggesting CoA was being recycled in the MCP. Such cofactor recycling could be employed with heterologous pathways as well.

Several single-enzyme encapsulation investigations demonstrate that the addition of a permeable barrier around an enzyme can sequester the product of an enzyme, suggesting that it could sequester the intermediate of a multi-enzyme pathway. For example, Patterson *et al.* encapsulated an alcohol dehydrogenase in the P22 virus-like particle and observed a change in the apparent enzyme kinetics in a permeability-dependent manner (31). One hypothesis was that the diffusion of the product was changing due to the presence of a diffusion barrier. Glasgow *et al.* also observed permeability-dependent changes to apparent enzyme kinetics by encapsulating an alkaline phosphatase in the MS2 VLP and mutating the pore residues to change the charge of the pore (32). Kinetic modeling suggested that product efflux was being reduced with certain mutants, demonstrating the sequestering of metabolites in the shell.

Here, we leveraged cell-free metabolic engineering approaches to determine whether encapsulation in MCPs can be used to favor the product of the encapsulated pathway over competing reactions. Our model system is the *Salmonella enterica* serovar Typhimurium LT2 Pdu MCP engineered to encapsulate the deoxyviolacein branch of the violacein biosynthesis pathway. We assessed pathway performance in a cell-free context, allowing us increased control over the reaction environment surrounding the Pdu MCP, as well as over the concentration of unencapsulated and encapsulated enzymes (33–35). We chose the violacein pathway for this work because it branches into four different colorimetric products: violacein (V), deoxyviolacein (DV), proviolacein (PV), and prodeoxyviolacein (PDV) (Figure 1A) (36, 37). We investigated the effects of pathway encapsulation on the violacein pathway enzymes by encapsulating a portion of the PDV branch. We observed a large reduction in net pathway flux, indicating that the introduction of a diffusion barrier may restrict enzyme access to its substrate. We next encapsulated a portion of the DV production branch and kept the V production branch not encapsulated. An increase in DV production and a decrease in V production was observed compared to when the DV branch and V branch were both not encapsulated. These results support the hypothesis that the Pdu MCP can divert pathway flux by providing a diffusion barrier between competing enzymes while colocalizing intermediates with the desired enzyme pathway components, a critical insight into how encapsulation could benefit both native metabolic pathways and heterologous pathways of interest.

**Figure 1:**
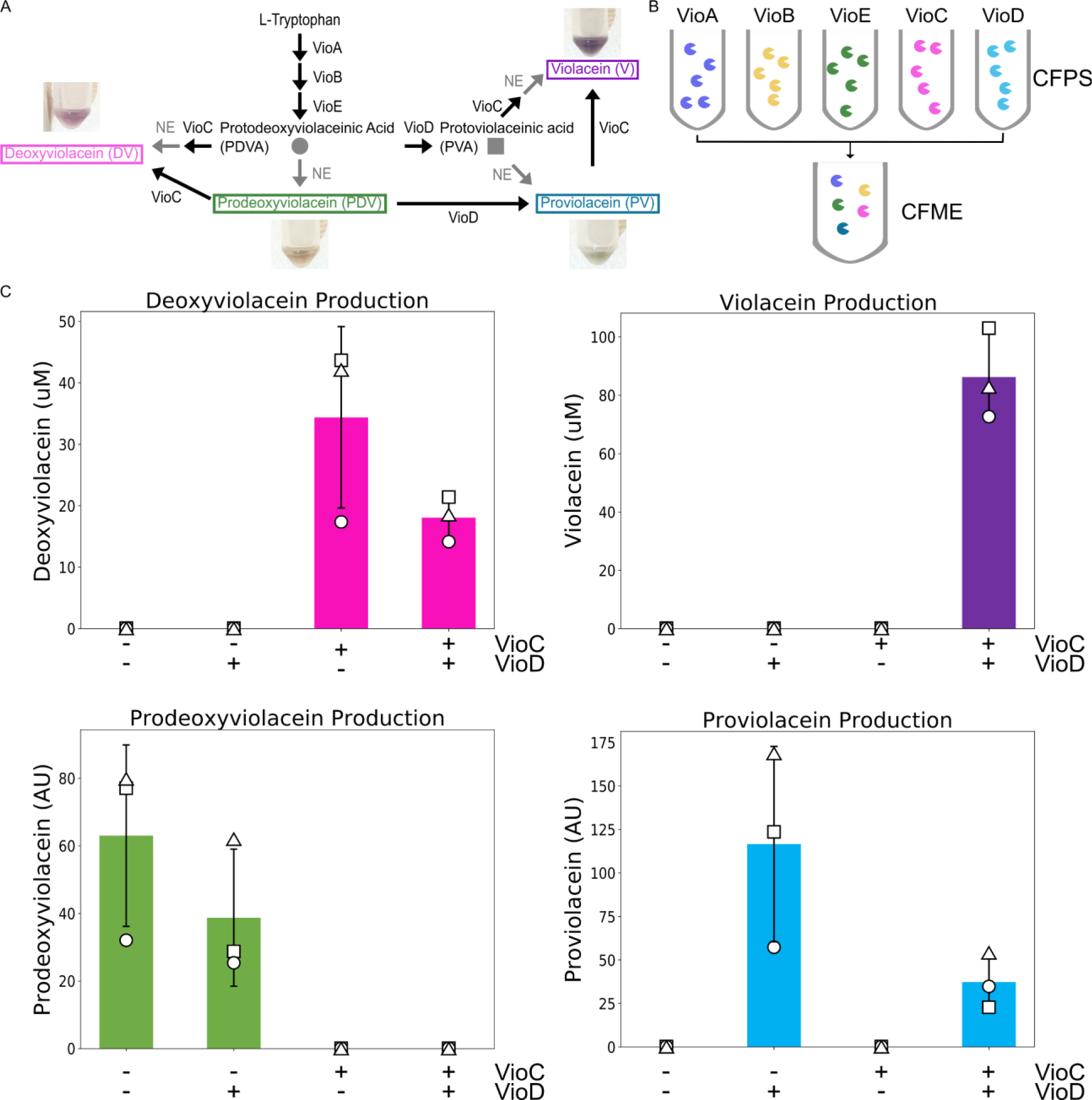
Production of violacein pathway products by CFME reactions -. A: Diagram of the violacein pathway highlighting the four main products: prodeoxyviolacein (PDV), deoxyviolacein (DV), proviolacein (PV), and violacein (V). Pictures taken after resuspending the CFME reactions in 100% methanol. B: Schematic of the process for making violacein pathway enzymes in CFPS and then combining them in CFME reactions to obtain metabolites. C: Production of violacein metabolites under different pathway enzyme combinations. Each CFME reaction includes VioA, VioB, and VioE and varies depending on whether VioC and/or VioD are present. Error bars denote +/- one standard deviation across technical replicates (N = 3). DV and V have commercial standards and are thus presented in molar concentrations. PV and PDV lack commercial standards, so we report quantities in arbitrary units (AU).

## Results

### The full violacein pathway is fully functional in CFME reactions

For our model metabolic pathway, we used the violacein pathway which converts tryptophan into V and the side products DV, PV, and PDV (Figure 1A). This pathway consists of the enzymes VioA, VioB, VioE, VioC, and VioD, and different combinations of these enzymes will determine which product is made. To gain control over the reaction environment and maintain equivalent enzyme concentrations across the samples and controls, we chose to study the pathway – in encapsulated and unencapsulated form - in a cell-free environment. Violacein is a known anti-microbial, so the cell-free system eliminates concerns about cellular fitness (37, 38). Nguyen *et al.,* Garamella *et al.,* and Pardee *et al.* each demonstrated violacein production in a cell-free system, but they used alternative cell-free backgrounds (39–41). Therefore, we first set out to confirm that we could produce the various pathway intermediates in our system. To do so, we used plasmid-based expression of each of the violacein pathway enzymes individually in cell-free protein synthesis (CFPS) reactions and then mixed the enzymes in cell-free metabolic engineering (CFME) reactions (Figure 1B). The quantity of full-length enzyme produced by each CFPS reaction was determined using ^14^C-leucine incorporation and autoradiograms (Table SI1 and Figure SI1) (42, 43), allowing us to hold constant the concentration of each violacein pathway enzyme across different experimental conditions in our CFME reactions. All CFME reactions were incubated for 20 hours before extracting the violacein metabolites and quantifying them.

We then combined the violacein enzymes in CFME reactions to determine if the violacein pathway was fully functional in the CFME context. When all of the violacein pathway enzymes are present in nature, the main product V and the side product DV are produced (Figure 1A) (36, 37). By controlling which violacein pathway enzymes were present in the CFME reaction, we were able to produce all four colorimetric metabolites (V, DV, PV, and PDV) (Figure 1C). For example, combining VioA, VioB and VioE results in PDV, while combining those three enzymes with VioC results in DV. We validated production of DV and V using high performance liquid chromatography (HPLC) because standards were commercially available for these two products. All of the products (V, DV, PV, and PDV) were confirmed via liquid chromatography mass spectrometry (LC-MS) and measured via HPLC (Figures S2 and S3).

### VioE and VioC tolerate Pdu MCP encapsulation peptides

In order to target the violacein pathway enzymes for encapsulation within the Pdu MCP, we needed to append encapsulation peptides to the enzymes. Encapsulation peptides are small peptides found on the N-termini of some of the native Pdu MCP enzymes (44). Fusing encapsulation peptides to heterologous proteins is sufficient to target the proteins for encapsulation within the Pdu MCP. To determine how much enzyme is present in MCPs produced and purified from cells, we also appended reporter tags for quantification. Thus, we had to assess the impact of adding these encapsulation peptides and reporter tags to the violacein pathway enzymes. We appended encapsulation peptides and reporter tags to the branching enzyme VioE and the enzyme required for DV production, VioC. To encapsulate VioE and VioC, we first fused the encapsulation peptide found on the N-terminus of PduD to VioE and VioC. We chose the PduD encapsulation peptide because it conferred the highest encapsulation efficiency of the known Pdu MCP encapsulation peptides for a model reporter protein (44). We fused a reporter FLAG tag to the C-termini of the VioE and VioC enzymes (Figure 2A) and PduD encapsulation peptide to the N-termini, connected by a GSGSGSG linker (for annotation purposes, ep denotes the N-terminal PduD encapsulation peptide and linker, and FL the C-terminal FLAG tag). We assessed the impact of tagging VioE by observing changes in PDV production in CFME reactions with VioA, VioB, and VioE^FL^ or epVioE^FL^. Similarly, we assessed the impact of tagging VioC by observing changes in DV production in CFME reactions with VioA, VioB, VioE, and VioC^FL^ or epVioC^FL^. The concentration of enzymes with and without encapsulation peptides was held constant between reactions. CFME reactions containing epVioE^FL^ and epVioC^FL^ produced less PDV and DV respectively than their counterparts without the encapsulation peptide. Specifically, the production of PDV in reactions with epVioE^FL^ was reduced by 33.15% compared to reactions with VioE^FL^ (*p* = 0.15) (Figure 2B), while DV production using reactions with epVioC^FL^ was reduced by 34.71% compared to those with VioC^FL^ (*p* = 0.044) (Figure 2C). Given the change in enzymatic activity when VioE^FL^ and VioC^FL^ are fused to the encapsulation peptide, for all remaining CFME experiments, we chose to use only the version of VioE and VioC with both the encapsulation peptide and FLAG tags present, whether or not they are encapsulated in MCPs. This will ensure that unencapsulated enzymes have the same activity as encapsulated enzymes, which must be fused to the encapsulation peptide to be encapsulated. Differences in pathway kinetics can therefore be attributed to enzyme encapsulation, without the confounding variable of changes in enzyme activity due to the presence or absence of any tag.

**Figure 2:**
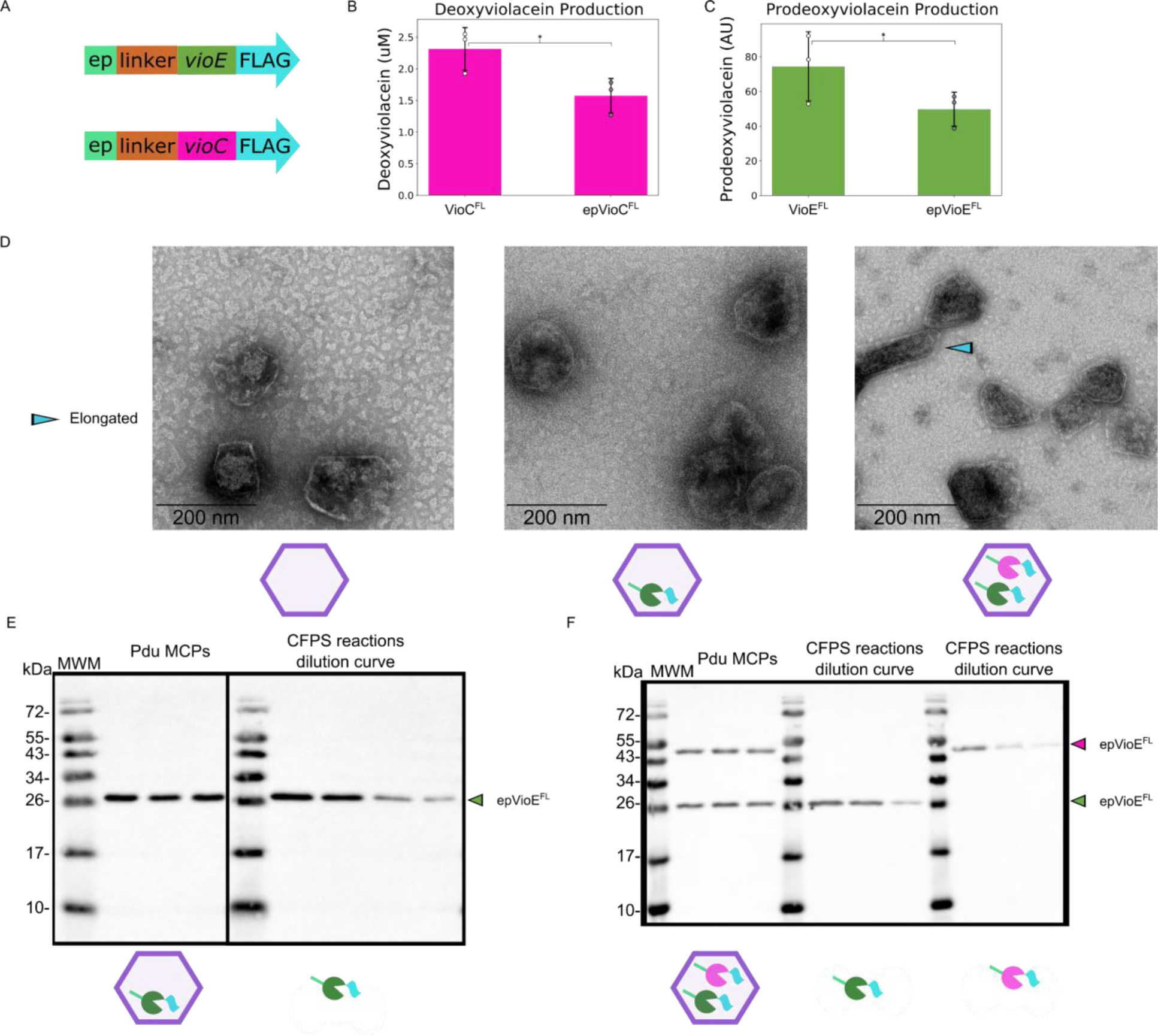
Characterization of VioE and VioC fused to signal sequences and encapsulated in the Pdu MCP –. A: Genetic constructs encoding the violacein enzymes N-terminally fused to the signal sequence of PduD (ep) with a GSGSGSG linker and C-terminally fused to FLAG tags (^FL^). The VioC^FL^ and VioE^FL^ enzymes are denoted with white-filled circles, and the epVioC^FL^ and epVioE^FL^ enzymes are denoted with grey filled circles. All CFME reactions have VioA and VioB added but not depicted. B: Production of PDV with VioE^FL^ and epVioE^FL^ in a CFME reaction. Error bars denote +/- one standard deviation from technical replicates (N = 3). Significance was determined with a two-tailed *t*-test and denoted with the following: * - *p* < 0.01, ** - *p* < 0.05, *** - *p* < 0.01, n.s. – *p* > 0.1. C: Production of DV with VioC^FL^ and epVioC^FL^ in a CFME reaction. Error bars and significance as previously described. D: Transmission electron micrographs of purified WT Pdu MCPs (left), Pdu MCPs with epVioE^FL^ encapsulated (middle), and Pdu MCPs with epVioE^FL^ and epVioC^FL^ encapsulated (right). Scale bars are 200 nm. E: Western blot of biological replicates (N = 3) of purified Δ*pduQ*::*epVioE^FL^* Pdu MCPs compared to a dilution series of epVioE^FL^ made in CFPS to confirm encapsulation. MWM, molecular weight marker. Spliced out lanes are indicated by the black borders. F: Western blot of biological replicates (N = 3) of purified Δ*pduD*::*epVioC^FL^*Δ*pduQ*::*epVioE^FL^* Pdu MCPs compared to dilution series of epVioE^FL^ and epVioC^FL^ made in CFPS to confirm encapsulation.

### VioE and VioC can be encapsulated in Pdu MCPs

The observed major product of the violacein pathway is V, but VioC is responsible for the conversion of an intermediate that leads to DV production if not further acted on by VioD. Therefore, we reasoned that encapsulating VioE and VioC together in the Pdu MCP and not encapsulating VioD would sequester the pathway intermediates to VioC and divert pathway flux to DV and away from V. To encapsulate VioE and VioC in the Pdu MCP, sequences encoding the epVioE^FL^ and epVioC^FL^ constructs were integrated into the *pduQ* and *pduD* loci of the *pdu* operon in the *Salmonella enterica* LT2 genome. These loci were chosen because of past successes in expression and encapsulation of a tagged fluorescent reporter with high efficiency (44, 45). Two strains were generated. In the first, the gene encoding epVioE^FL^ was inserted into the *pduQ* locus (Δ*pduQ*::*epVioE^FL^*). Within the second strain, genes encoding epVioE^FL^ and epVioC^FL^ were integrated at the *pduQ* and *pduD* loci (Δ*pduD*::*epVioC^FL^*Δ*pduQ*::*epVioE^FL^*). Pdu MCPs purified from these strains were run on SDS PAGE gels that were subsequently stained with Coomassie blue (Figure SI4). Bands were detected that were at the expected sizes for Pdu MCP proteins, suggesting that Pdu MCPs were purified. To verify that epVioE^FL^ and epVioC^FL^ were encapsulated in the Pdu MCP, we performed western blots against the FLAG tags. Successful detection of the FLAG tags suggested that the enzymes are encapsulated and co-purified with the Pdu MCPs. Western blots on purified Pdu MCPs from Δ*pduQ*::*epVioE^FL^* revealed bands at the expected size of ssPduD::VioE^FL^, indicating successful encapsulation in the Pdu MCP (Figure 2E). Similarly, western blots on purified Pdu MCPs from Δ*pduD*::*epVioC^FL^*Δ*pduQ*::*epVioE^FL^* revealed bands at the sizes expected for epVioE^FL^ and epVioC^FL^, providing strong supportive evidence that both enzymes are encapsulated in the Pdu MCP (Figure 2F).

We also wanted to determine if the genomic modification and encapsulation of heterologous enzymes affected the microcompartment assembly. Pdu MCPs were purified from the mutants and imaged by transmission electron microscopy (TEM) to identify possible changes to the Pdu MCP structure (Figure 2D). TEM revealed the Pdu MCPs from the Δ*pduQ::epVioE^FL^* strain share similar morphology to wildtype (WT) Pdu MCPs. The Pdu MCPs from the Δ*pduD::epVioC^FL^* Δ*pduQ::epVioE^FL^* strain were also mostly similar in morphology to WT MCPs, although some were elongated. This may occur because modifications to the *pdu* operon that have polar effects on the expression levels of shell proteins, particularly the vertex protein PduN, can lead to the formation of elongated structures (24, 46).

We next determined the concentration of enzymes encapsulated within the Pdu MCP, so that in control CFME reactions we could add the same amount of enzyme as was present in our MCP-containing CFME reactions. Western blotting was performed against the FLAG tag on Pdu MCPs purified from the strain Δ*pduQ*::*epVioE^FL^* as well as on a sample of the same epVioE^FL^ made by CFPS (Figure 2E). The concentration of enzymes produced in the CFPS reaction was quantified by ^14^C- leucine incorporation and autoradiograms, allowing us to use densitometry to calculate the concentration of each encapsulated enzyme. We conducted the same approach for Pdu MCPs purified from the strain Δ*pduD*::*epVioC^FL^* Δ*pduQ*::*epVioE^FL^* and the epVioE^FL^ and epVioC^FL^ made by CFPS (Figure 2F). The concentration of enzymes in CFME reactions was normalized to the amount of epVioE^FL^ encapsulated in Pdu MCPs purified from the Δ*pduD*::*epVioC^FL^* Δ*pduQ*::*epVioE^FL^* strain. Thus the amount of Pdu MCPs purified from Δ*pduQ*::*epVioE^FL^* strains added to CFME reactions was adjusted to match the concentration of epVioE^FL^. We then matched the concentration of enzymes in the control CFME reactions by adding CFPS generated enzymes to ensure the total concentration of each enzyme in the sample was held constant across the unencapsulated and encapsulated enzyme conditions.

### Encapsulation of epVioE^FL^ reduces overall PDV production

We hypothesize that encapsulation of heterologous pathways in the Pdu MCP may divert and enhance the flux of the pathways by providing a diffusion barrier around pathway enzymes and metabolites, but this same barrier may also impact substrate access to the first encapsulated enzyme (21, 22). To that end, when epVioE^FL^ is encapsulated, the diffusion barrier of the MCP shell may restrict epVioE^FL^’s access to its substrate, an indole-3-pyruvic acid (IPA) imine dimer, resulting in reduced production of PDV. To investigate the effects of introducing a diffusion barrier between epVioE^FL^ and the IPA imine dimer, we conducted CFME reactions with VioA and VioB produced by CFPS and epVioE^FL^, the latter of which was sourced either from a CFPS reaction or from purification of Pdu MCPs produced by bacteria expressing that protein (Figure 3). The concentration of epVioE^FL^ was held constant between reactions with and without MCPs. CFME reactions with epVioE^FL^ encapsulated produced 96.6% less PDV compared to CFME reactions with unencapsulated epVioE^FL^ (*p =* 0.0057). While there was a large reduction in PDV production, some PDV was still produced, suggesting that the IPA imine dimer is still able to diffuse into the Pdu MCP. It also demonstrates that the epVioE^FL^ enzyme is still functional in the Pdu MCP. The reduction in PDV production likely resulted from the diffusion barrier and/or a potential change in the apparent enzyme kinetics of epVioE^FL^ when it is encapsulated.

**Figure 3:**
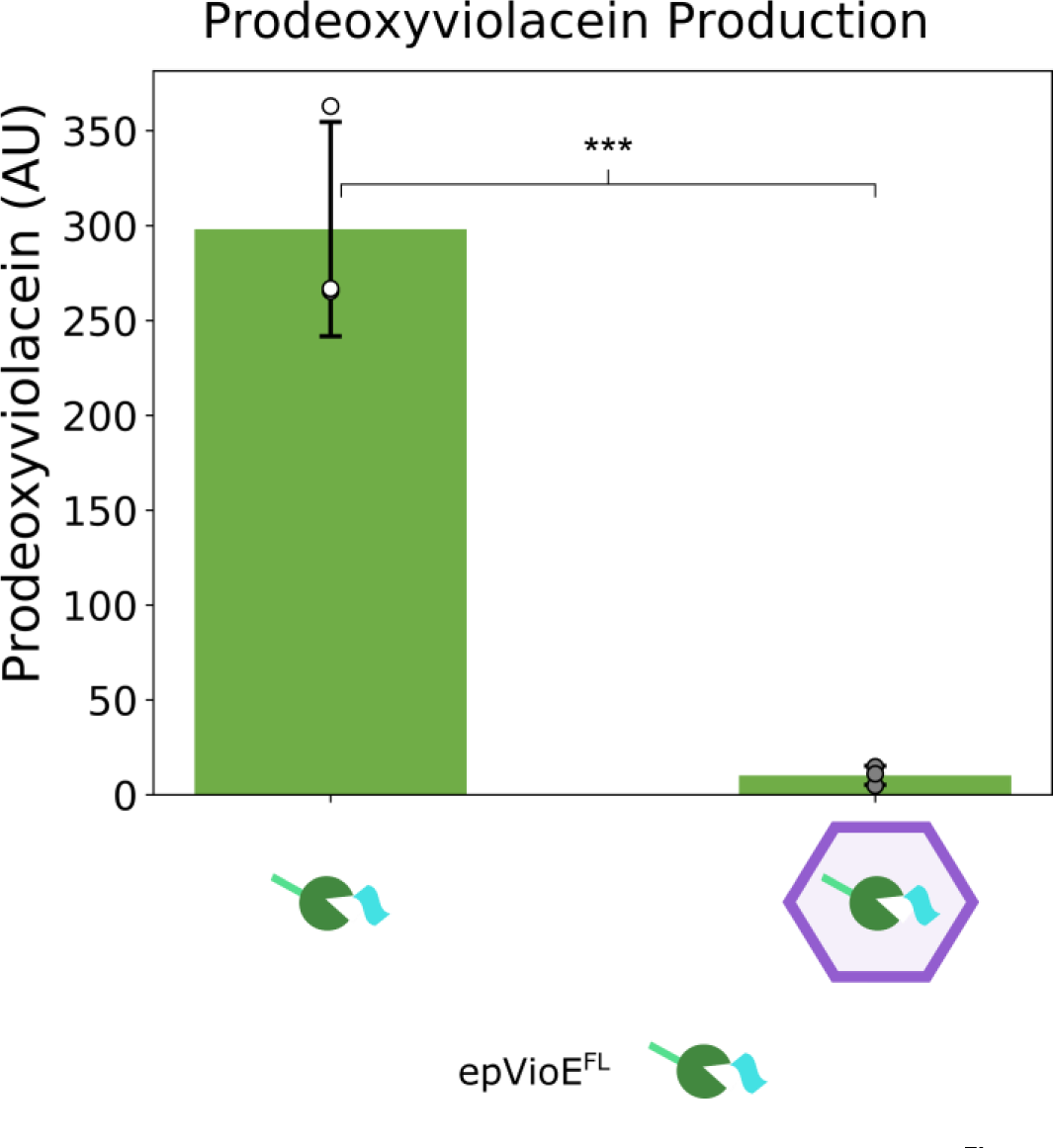
PDV production when epVioE^FL^ is encapsulated in the Pdu MCP. Production of PDV in CFME reactions with unencapsulated or encapsulated epVioE^FL^. VioA and VioB are added to the CFME reaction but are not depicted. Error bars denote +/- one standard deviation from technical replicates (N = 3) for the unencapsulated epVioE^FL^ condition or biological replicates (N = 3) for the encapsulated epVioE^FL^ condition. The unencapsulated epVioE^FL^ condition is denoted with white-filled symbols, and encapsulated epVioE^FL^ condition is denoted with grey filled circles. Significance was determined with a two-tailed *t*-test and is denoted by the following: * - *p* < 0.01, ** - *p* < 0.05, *** - *p* < 0.01, n.s. – *p* > 0.1.

### Encapsulation of epVioC^FL^ does not influence DV production

To test our hypothesis using the encapsulated DV branch, epVioC^FL^ needs to be encapsulated with epVioE^FL^ in the Pdu MCP. To this end, we first needed to establish that encapsulating epVioC^FL^ would not change the overall production of DV by the end of the assay. Otherwise, changes to DV and V production may be attributed to epVioC^FL^ not tolerating encapsulation rather than the introduction of diffusion barriers to sequestering pathway intermediates to epVioC^FL^. Therefore, we conducted CFME reactions in which epVioE^FL^ was encapsulated, while epVioC^FL^ was either encapsulated or not (Figure 4A). All enzyme concentrations were held constant across conditions. There was no significant difference in the amount of DV produced between CFME reactions with epVioC^FL^ encapsulated compared with not encapsulated (*p* = 0.84). Since VioC can convert both PDVA and PDV into DV, it was expected that no PDV would be detected by the end of the reaction. From this data, we determined that epVioC^FL^ can tolerate encapsulation within the Pdu MCP with no significant observable change to DV production.

**Figure 4.**
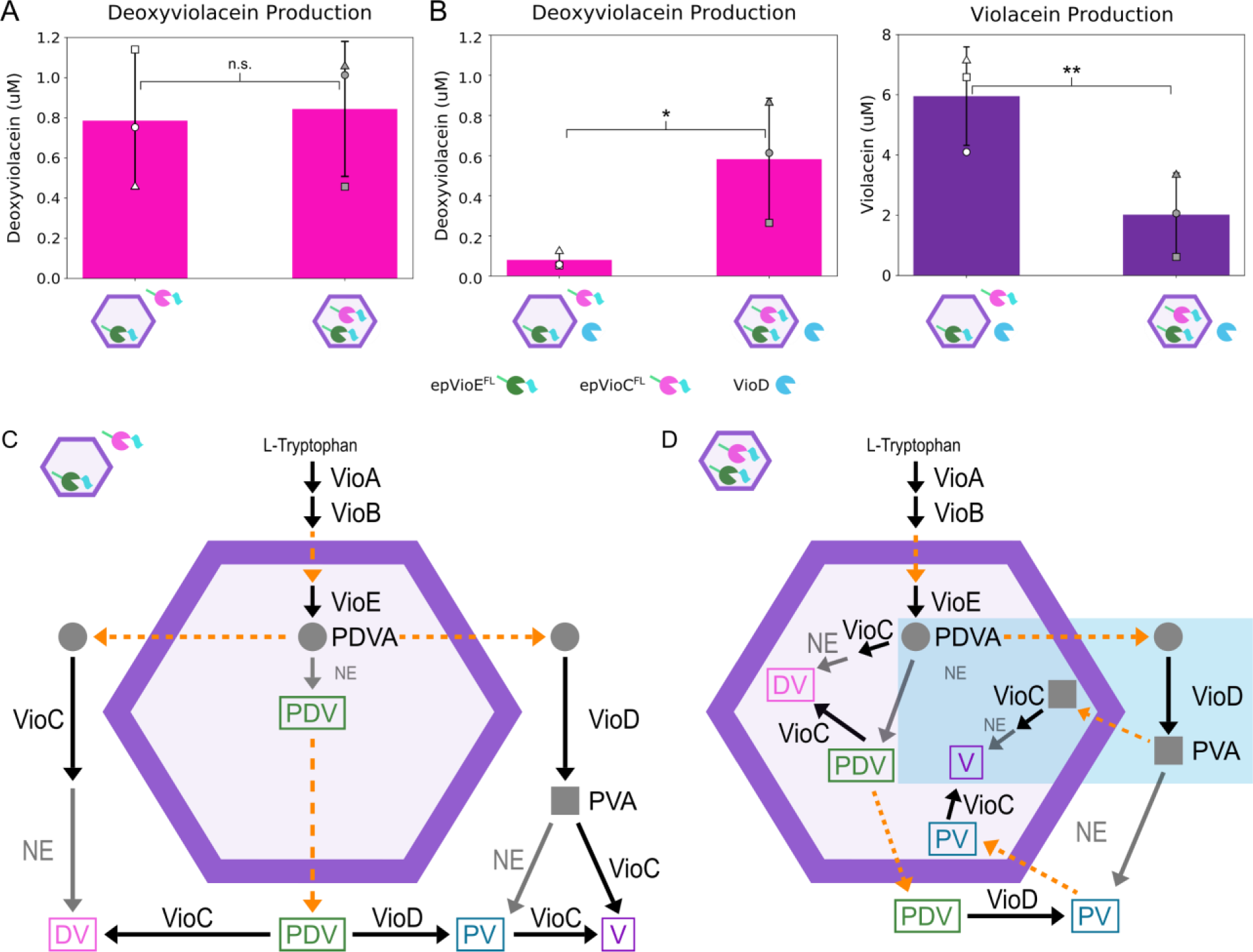
Production of violacein pathway products deoxyviolacein and violacein using various enzyme encapsulation schemes. All CFME reactions have VioA and VioB added but not depicted. Error bars denote +/- one standard deviation from biological replicates (N = 3). The encapsulated epVioE^FL^ and not encapsulated epVioC^FL^ condition is denoted with white-filled symbols, and the encapsulated epVioE^FL^ and epVioC^FL^ condition is denoted with grey filled symbols. The first replicate is represented by a □, the second replicate is represented by a Δ, and the third replicate is represented by a O. Significance was determined with a two tailed *t*-test and is denoted with the following: * - p < 0.01, ** - p < 0.05, *** - p < 0.01, n.s. – p > 0.1. A: Production of DV in CFME reactions with Pdu MCPs with epVioE^FL^ encapsulated and epVioC^FL^ not encapsulated or with Pdu MCPs with epVioE^FL^ and epVioC^FL^ encapsulated. B: Production of DV and V in CFME reactions with VioD not encapsulated and with Pdu MCPs with epVioE^FL^ encapsulated and epVioC^FL^ not encapsulated or with Pdu MCPs with epVioE^FL^ and epVioC^FL^ encapsulated. C: Diagram of the production of PDVA (grey circle), PVA (grey square), V, DV, PV, and PDV when epVioE^FL^ is encapsulated and epVioC^FL^ and VioD are not encapsulated. Black arrows represent enzymatic reaction, grey arrows indicate non- enzymatic reaction, and dashed orange arrows indicate diffusion across the Pdu MCP shell. D: Diagrams of the production of PDVA (grey circle), PVA (grey square), V, DV, PV, and PDV when epVioC^FL^ and epVioE^FL^ are encapsulated and VioD is not encapsulated. An example pathway to produce V is highlighted in a light blue box.

### Introducing diffusion barriers between the encapsulated pathway and competing enzymes diverts pathway flux to the encapsulated pathway

We next determined if intermediate retention could divert pathway flux towards the encapsulated enzymes and away from competing enzymes. VioD interacts with either the PDVA intermediate or PDV to produce PV which can then be converted into V by VioC and a non-enzymatic reaction. If VioC interacts with PDVA or PDV first, DV is produced instead. We hypothesized that co- encapsulating epVioE^FL^ and epVioC^FL^ would prevent VioD from interacting with the PDVA or PDV, thus increasing DV production while decreasing V production. In addition to providing a diffusion barrier, encapsulation also co-localizes VioE and VioC, increasing the local PDVA and PDV concentration in proximity to VioC, whereas VioD is not colocalized. The benefit of co-localizing VioE and VioC was highlighted by Zhao *et al.* using the optoDroplet and PixELL systems to cluster VioE and VioC together in a liquid-like droplet, which enhanced both DV production and specificity (47).

To test our hypothesis, we conducted CFME reactions in which epVioE^FL^ and epVioC^FL^ are encapsulated inside the Pdu MCP while VioD remains unencapsulated. The concentration of VioA, VioB, epVioE^FL^, epVioC^FL^, and VioD in each reaction was held constant. A key control was encapsulating epVioE^FL^ alone while epVioC^FL^ and VioD are not encapsulated as it provides the metabolite profile of non-diverted pathway flux (Figure 4C). Specifically, when epVioE^FL^ is encapsulated alone, PDVA and PDV are produced and then trapped in the Pdu MCP. As PDVA and PDV diffuse out, VioD and epVioC^FL^ have equal access to them to produce DV and V, similar to the scenario for the native unencapsulated pathway. For this case, we see that DV production is an order of magnitude lower for CFME reactions when epVioE^FL^ is encapsulated alone and VioD is present compared to the same CFME reactions without VioD (*p* = 0.067) (Figures 4A and 4B). This result highlights how much DV production is reduced when the competing VioD enzyme is present. It is not surprising, considering that in nature V is the primary product of the violacein pathway and DV is produced as a side product. As we hypothesize above, we expect that when epVioE^FL^ and epVioC^FL^ are co-encapsulated and VioD is not encapsulated, PDVA and PDV are likely trapped with epVioC^FL^ for an extended time. This allows epVioC^FL^ to act on PDVA and PDV to produce DV before they diffuse out and are acted upon by VioD. Indeed, with these CFME reactions, we observed a significant increase in DV production and a significant reduction in V production, as compared to when epVioC^FL^ was left unencapsulated with VioD (DV: *p* = 0.084; V: *p* = 0.034) (Figure 4B). Thus, pathway flux was diverted to DV when both epVioE^FL^ and epVioC^FL^ were encapsulated. We note that there are two additional diffusion barriers for V production (compared to DV production) in this scheme regardless of the path taken to that product (Figure 4D). For example, the PDVA intermediate would need to diffuse out of the MCP to be converted into PVA by VioD, then PVA would need to diffuse back into the MCP to be converted by epVioC^FL^ and a non-enzymatic reaction into V (Figure 4D, blue box). As expected, PDV was not detected under either condition since PDV would be converted either by epVioC^FL^ and a non-enzymatic reaction into DV, or by VioD into PV (36). PV production was not significantly different between CFME reactions with encapsulated and unencapsulated epVioC^FL^ (PV: *p* = 0.92) (Figure SI5). PV can be converted by epVioC^FL^ and a non-enzymatic reaction into V, so any PV would eventually be converted to V regardless of epVioC^FL^’s encapsulation status. The lack of change we observed in PV production implies that the lower V production is not due to net pathway flux being diverted to PV. Overall, these results demonstrate that encapsulation within the Pdu MCP can divert violacein pathway flux by introducing diffusion barriers between the pathway intermediates and external competing enzymes.

## Discussion

This is the first report to demonstrate the long-hypothesized idea that encapsulation in MCPs influences the distribution of pathway end-products. When epVioE^FL^ and epVioC^FL^ were encapsulated in the Pdu MCP in CFME reactions with unencapsulated VioD, the product profile shifted towards DV and away from V compared to the unencapsulated epVioC^FL^ control. This work demonstrates the potential of Pdu MCPs to expand the metabolic engineering toolkit and supports our hypothesis that bacteria utilize MCPs for product selectivity.

This is also the first report of employing CFME to study the impact of pathway encapsulation in MCPs, which required substantial methods development. Prior studies on the violacein pathway in a cell-free context demonstrated the importance of controlling the concentration of each protein added to the CFME reaction, which they did by tuning plasmid concentration in the corresponding CFPS reactions (39–41). We improved this procedure by instead holding constant the concentration of each protein added to the CFME, as determined by assessing the concentration of full protein made in each CFPS reaction and present in each purified MCP sample. This ensured that the primary difference between conditions was whether the enzymes were encapsulated or not. Likewise, prior to this work, encapsulated protein in the Pdu MCP was measured semi-quantitatively with fluorescence and densitometry of western blots (44, 48, 49). The use of ^14^C-leucine incorporation and autoradiograms allowed us to create protein standards that we could run on western blots to compare to the same encapsulated enzyme constructs encapsulated in the Pdu MCP using densitometry. To date, this is the first fully quantitative measurement of heterologous protein encapsulation in the Pdu MCP. Using CFPS to generate standards for western blotting is quicker and easier than purifying proteins, especially those that are challenging to express *in vivo* and purify.

Another advantage of using a cell-free system over an *in vivo* one was that we could compare metabolic profiles across scenarios in which pathway enzymes are and are not encapsulated. *In vivo,* it is difficult to control the percentage of encapsulation-tagged proteins present in the MCP versus the cytosol. For example, to ensure we had unencapsulated enzymes present *in vivo*, we would have had to remove the encapsulation peptide, which we show *in vitro* with the violacein pathway enzymes can lead to differences in enzyme kinetics.

The unencapsulated enzymes in our experiments were made from CFPS reactions whereas encapsulated enzymes in the Pdu MCPs were purified from bacterial cultures. With our analysis, we assume that these differences in enzyme preparation are not responsible for observed differences across the unencapsulated and encapsulated conditions. This assumption is supported by our results showing DV production is unchanged whether epVioC^FL^ is encapsulated (biologically produced, in MCPs) or unencapsulated (CFPS produced, Figure 4A).

According to our results, the Pdu MCP can be used to favor the production of the encapsulated pathway over competing pathways. When epVioE^FL^ was encapsulated and both epVioC^FL^ and VioD were unencapsulated, the production of V was favored. When epVioE^FL^ and epVioC^FL^ were both encapsulated and VioD was on the outside of the Pdu MCP, the product distribution favored the DV branch instead of the V branch. We attribute this behavior to the addition of two diffusion barriers in the production of V as well as the removal of a diffusion barrier between pathway intermediates PDVA and PDV and epVioC^FL^ (Figure 4C and 4D). It is important to note that we cannot close the mass balance for the diversion of net pathway flux from V to DV as the increase in DV does not account fully for the decrease in V. A limitation of this study is the inability to measure all of the intermediates of the violacein pathway, such as the intermediate product of VioC before it non-enzymatically converts to DV. Some of the decrease in V production could be trapped in intermediates we cannot measure.

The diversion of pathway flux with selective enzyme encapsulation may not be generalizable to all metabolic pathways. According to kinetic models in Jakobson *et al*., there is a predicted optimal Pdu MCP shell permeability for the 1,2-propanediol utilization pathway that maximizes the concentration of intermediate in the Pdu MCP to promote pathway performance. If shell permeability to the pathway metabolites is too low, then the substrate will not be able to diffuse in and be acted upon by the encapsulated enzymes. If the shell permeability is too high, the intermediate will leak out as it is being produced, resulting in no intermediate being sequestered to the encapsulated enzymes. The optimal shell permeability would allow sufficient substrate diffusion into the MCP while still sequestering the intermediate inside. If we assume the permeability of the Pdu MCP has evolved to be close to optimal for increasing flux through the Pdu pathway, then we expect the permeability to be lower for the larger violacein metabolites (19, 21). The IPA imine dimer (∼404.42 g/mol) is much larger than the native Pdu MCP substrate, 1,2-PD (76.09 g/mol). Indeed, there was a stark reduction in PDV production when epVioE^FL^ is encapsulated in the Pdu MCP, suggesting the IPA imine dimer cannot diffuse easily in the Pdu MCP. To ameliorate this issue, one could mutate the pores of the Pdu MCP shell to achieve a permeability nearer to the optimum that allows for selectivity at higher overall yields (25, 50–53). While permeability is one factor, the reaction rates of the pathway steps are also important. In Jakobson *et al.,* kinetic modeling revealed that the degree to which three spatial organization techniques (encapsulation in an organelle, colocalization on open scaffolds, or no organization whatsoever) impacted pathway performance depending on enzyme kinetics in the pathway as well as shell permeability to the pathway metabolites (22). For some pathways, open scaffolds can confer significant changes to observed pathway kinetics without decreasing yield due to permeability limits (54–57). Characterizing the permeability of the Pdu MCP to target substrates, intermediates, and products will allow us to utilize such kinetic models to choose which pathways would most benefit from MCP encapsulation versus colocalization on an open scaffold.

Taken together, our work shows that intermediate sequestration can be a benefit of pathway encapsulation in MCPs and highlights the potential of heterologous pathway encapsulation in the Pdu MCP as a metabolic engineering tool for pathway flux diversion, both *in vivo* and *in vitro*.

## Methods and Materials

### Bacterial strains and plasmids

*Escherichia coli* BL21 DE3 was used for the preparation of cell extracts, which were used to generate pathway enzymes and to conduct reactions with those enzymes (33, 58). The genes for the violacein pathway enzymes were generously provided by the John Dueber lab on one plasmid. Sequences encoding the violacein enzymes were cloned via Gibson cloning into a pJL1 parent vector (Addgene, 69496) (59). Gibson cloning was also used to fuse the violacein enzymes with an N-terminal ssPduD tag and a GSGSGSG glycine-serine linker and a C-terminal FLAG tag. All primers used to amplify inserts for Gibson cloning can be found in (Table SI2). All plasmids used in this study and their key characteristics can be found in (Table SI2).

*Salmonella enterica* serovar Typhimurium LT2 was used to purify 1,2-propanediol utilization microcompartments (Pdu MCPs) encapsulating the violacein pathway enzymes.

### Recombineering

Genomic modifications to *Salmonella enterica* serovar Typhimurium LT2 (LT2) strains were introduced using the λ red recombineering technique as previously described (60). Briefly, LT2 strains were transformed with the plasmid pSIM6 which contains the λ red recombineering machinery as well as carbenicillin resistance. The plasmid pSIM6 expresses the λ red recombineering machinery at 42 °C and is ejected from the cell at 37 °C. LT2 with pSIM6 was transformed with DNA products using electroporation and then LT2 was recovered at 30 °C, to retain the pSIM6 plasmid, or 37 °C, to remove the pSIM6 plasmid. At the locus of interest, we inserted a cassette containing a chloramphenicol resistance gene (*cat*) and a sucrose sensitivity gene (*sacB*) amplified from the TUC01 genome. Cassette incorporation was confirmed by growing cells on lysogeny broth (LB)-Agar supplemented with 10 μg/mL of chloramphenicol and 30 μg/mL of carbenicillin. Single colonies that grew on the chloramphenicol plates were streaked onto 6% (w/w) sucrose plates to assess sucrose sensitivity. The *cat/sacB* cassette was then replaced with a PCR product of a full gene with 50 base pairs homology upstream and downstream of the locus of interest. Incorporation of the DNA of the desired gene was screened with sucrose sensitivity; polymerase chain reaction was conducted on clones at the locus of interest to be confirmed with Sanger sequencing (Genewiz).

### Pdu MCP purification

Pdu MCPs were purified using a differential centrifugation method as previously described (61, 62). Briefly, to purify Pdu MCPs, we inoculated single colonies of LT2 strains LB liquid media and grew them at 30 °C for 24 hours. The overnight culture was subcultured 1:1,000 into 200 mL of No Carbon Essential (NCE) media supplemented with 50 µM ferric citrate, 1 mM magnesium sulfate, 42 mM succinate as a carbon source, and 55 mM 1,2-propanediol. The NCE subculture was grown at 37 °C at 225 rpm to an OD600 of 1-1.5. Cells were harvested via centrifugation and then resuspended in lysis buffer. Resuspended cells were incubated in this lysis buffer at room temperature for 30 minutes. After lysis, the lysate was kept on ice for 5 minutes. The lysate was then clarified using centrifugation. To separate the MCPs from the clarified lysate, they were spun down via centrifugation. The supernatant was removed, and the MCP pellet was washed with buffer (32 mM Tris-HCl, 200 mM KCl, 5 mM MgCl2, 0.6% (v/v) 1,2-propanediol, 0.6% (w/w) octylthioglucoside, pH 7.5–8.0). MCPs were pelleted, and the supernatant was removed. The MCP pellet was resuspended in buffer B (50 mM Tris-HCl, 50 mM KCl, 5 mM MgCl2, pH 8.0) and stored at 4 °C until use. The concentration of the purified Pdu MCPs was determined using a bicinchoninic acid assay (Thermo Scientific 23225).

### Protein gels with Coomassie staining

Purified Pdu MCPs were diluted to a concentration of 112.5 µg/mL in 2 M Urea and 1x Laemmli buffer (62.5 mM Tris pH 6.8, 2% sodium dodecyl sulfate (SDS), 10% glycerol, 0.05% bromophenol blue) with 10% β-mercaptoethanol. The diluted sample was boiled at 95°C for 15 minutes and frozen at -20 °C for storage. 1.5 µg of boiled Pdu MCPs were loaded on a 15% SDS-polyacrylamide gel electrophoresis (SDS-PAGE) gel and run at 120 V for 100 minutes. Afterward, the gel was stained with Coomassie Brilliant Blue R-250 (IGN Biomedicals Inc 04-821- 616) and imaged on an Azure 600 imaging system.

### Cell extract preparation

The *E. coli* BL21 DE3 (Life Technologies) cells were prepared into cell-free extracts via growth, harvest, lysis, and preparation as previously described (58, 63). Briefly, cells were cultured in full-baffled flasks with 1 L of 2xYTPG (16 g/L tryptone, 10 g/L yeast extract, 5 g/L NaCl, 7 g/L potassium phosphate monobasic, 3 g/L potassium phosphate dibasic, 18 g/L glucose) at 37 °C. Once the cultures reached an OD600 of 0.4, 1 mM IPTG was added to induce expression of the T7 polymerase. At an OD600 of 0.4-0.6, cells were harvested via centrifugation at 5,000 x g for 10 minutes at 4 °C. The supernatant was removed from the cells and the remaining pellets were washed three times with cold S30 buffer (10 mM tris acetate, pH 8.2, 14 mM magnesium acetate, and 60 mM potassium acetate). Each wash was followed by centrifugation at 10,000 x g at 4 °C. After the last wash, cells were flash frozen with liquid nitrogen and stored at -80 °C.

Before lysing the cells, cell pellets were thawed and resuspended in 1 mL of cold S30 buffer. The resuspended cells were then lysed using an EmulsiFlex-B15 homogenizer (Avestin) in a single pass at a pressure of 20,000-25,000 psi. After, cell debris was removed by spinning the cells at 12,000 x g for 30 minutes at 4 °C. Supernatant was moved to subsequent tubes to be flash frozen in liquid nitrogen and stored at -80 °C until use.

### CFPS

CFPS was conducted according to methods previously described (64, 65). Briefly, 15.6 µL CFPS reactions set up to produce proteins for use in cell-free metabolic engineering (CFME). The 15.6 µL CFPS reactions contained the following: 6 mM Mg(Glu)2, 10 mM NH4(Glu), 130 mM K(Glu), 1.2 mM ATP, 0.85 mM GTP, 0.85 mM UTP, 0.85 mM CTP, 0.034 mg/mL folinic acid, 0.171 mg/mL tRNA, 33.33 mM PEP, 2 mM 20 standard amino acids, 0.33 mM NAD+, 0.27 mM CoA, 4 mM CoA, 1 mM putrescine, 1.5 mM spermidine, and 57 mM HEPES. To start the CFPS reaction, 200 ng of pJL1 plasmid containing our enzymes of interest was added to the reaction mixture. The CFPS reactions were pipetted into 2 mL microcentrifuge tubes (Fisher Scientific Cat# 02-682-004) and incubated at 30 °C for 20 hours. In addition to producing our enzymes of interest, super folder GFP was made in three technical replicates to determine if there was high variability in the CFPS reactions. To measure GFP fluorescence, the completed reactions were diluted 1:50 in MilliQ water and measured on a Biotek Synergy H1 plate reader. If the coefficient of error exceeded 20% for the GFP fluorescence, then the CFPS reactions from that batch were not used.

### 14C-leucine incorporation

Enzyme concentration was quantified using ^14^C-leucine incorporation during the CFPS reaction as previously described (43). 10 µM of ^14^C-leucine was added to the CFPS reaction along with all 20 standard amino acids. The CFPS reactions were stopped by the addition of an equal volume of 5 N KOH. Afterward, total and soluble fractions of the CFPS reaction were collected, respectively. Using 10% trichloroacetic acid, we precipitated out the proteins. Radioactive counts were measured from the precipitated proteins using liquid scintillation (PerkinElmer MicroBeta^2^ 2450 Microplate Counter) to get quantitative measurements of the soluble and total yields of the enzymes produced in CFPS. All enzyme concentrations from CFPS are listed in Table SI1 (43, 64).

### Autoradiogram

In addition to using ^14^C-leucine incorporation to quantify protein concentration from CFPS reactions, autoradiography was also used to measure how much ^14^C-leucine was incorporated into complete protein versus truncated protein products. We ran ^14^C-leucine incorporated CFPS reactions on an SDS-PAGE gel and then exposed the gels using autoradiography. A Typhoon 7000 (GE Healthcare Life Sciences) was used to image the autoradiograms. Using ImageJ, we performed densitometry on the autoradiogram to determine how much ^14^C-leucine was incorporated into the complete protein by comparing the densitometry of the band that matched the size of complete protein to the densitometry of all bands in the autoradiogram.

### Western blotting and densitometry

SDS-PAGE gels for western blotting were loaded with Pdu MCPs using the same concentrations as the SDS-PAGE gels for Coomassie staining and were run at 150 V for 1 hour. Samples were transferred to a polyvinylidene fluoride (PVDF) membrane made specifically for fluorescent secondary antibodies (Immobilon IPFL00010) using a Bio-Rad Transblot SD at 25 V, 150 mA, for 35 min. The membrane was blocked using tris-buffered saline with 0.1% Tween® 20 detergent (20 mM Tris, 150 mM sodium chloride, 0.05% (v/v) Tween 20, pH 7.5) with 5% bovine serum albumin (w/w) for 1 hour at room temperature. Mouse anti-FLAG M2 antibody (Millipore Sigma F3165) was then applied to the membrane diluted 1:6,666 in 1% (w/w) bovine serum albumin for 1 hour. Anti-FLAG antibody was removed, and the membrane was washed four times with 0.05% Tween® 20 detergent. Afterwards, goat anti-mouse IgG polyclonal antibody (IRDye® 680RD) (Licor) was applied to the membrane for 1 hour at room temperature and then washed four times with 0.05% Tween® 20 detergent. The membrane was then imaged on an Azure 600 (680 nm excitation,694 nm emission) for fluorescent secondary antibody. Densitometry on the western blot was conducted with ImageJ.

### Determining how much violacein enzyme was encapsulated in the Pdu MCP

Densitometry of epVioE^FL^ and epVioC^FL^ made from CFPS and encapsulated in Pdu MCPs was performed using ImageJ software. For epVioE^FL^ and epVioC^FL^ made using CFPS, moles of protein were measured using ^14^C-leucine incorporation and then normalized by the densitometry of the autoradiogram (Equation 1). We determined moles of protein per AU of densitometry on the western blot for epVioE^FL^ and epVioC^FL^ by plotting densitometry against moles of protein (Equation 2). In Equation 2, f is the slope of the linearization of moles of protein versus AU. The moles of each enzyme encapsulated in the Pdu MCPs were measured and used to normalize the concentration of enzyme in each CFME condition across the unencapsulated and encapsulated controls.

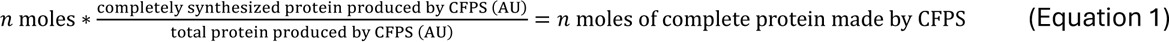

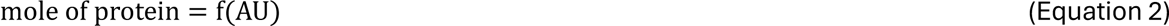

### Transmission electron microscopy

For transmission electron microscopy, 400-mesh copper grids with Formvar/carbon film (EMS Cat# FCF400-Cu-50) were hydrophilized using a glow discharge system. A 10 µL aliquot of purified Pdu MCP sample was deposited onto the grid. After 5–10 seconds, the sample was wicked away, followed by the application of 10 µL of a 1% (w/v) aqueous uranyl acetate (UA) solution as a negative stain. The UA was immediately wicked away, and the process was repeated with two additional UA applications: the first was wicked away immediately, and the second was allowed to sit for 4 minutes before removal. The grids were then imaged using a JEOL 1400 Flash transmission electron microscope equipped with a Gatan OneView Camera at 50,000 x magnification at room temperature. Images were cropped using ImageJ.

### Statistical methods and individual data point representation

For figures where significant differences were being investigated, significance was determined with a two tailed *t*-test and is denoted with the following: * - p < 0.1, ** - p < 0.05, *** - p < 0.01, n.s. – p > 0.1. For CFME reactions with no Pdu MCPs, the reactions were done in technical replicates (N = 3). For reactions with Pdu MCPs included, biological replicates (N = 3). All of the bar graphs are presented as mean +/- one standard deviation. The metabolite measurements of individual replicates are juxtaposed on the bar graphs. For CFME reactions where more than one metabolite is being measured, three different symbols were used to show the first, second, and third replicate, so the reader can track what replicate corresponds to each metabolite profiles. The first replicate is represented by a □, the second replicate is represented by a Δ, and the third replicate is represented by a O.

### CFME

We generated 30 µL cell-free metabolic engineering (CFME) reactions by mixing proteins synthesized in CFPS reactions and purified Pdu MCPs. The concentration of each violacein pathway enzyme in the CFME reactions was held constant across the experimental conditions within an experiment. For experiments where we compared conditions with Pdu MCPs with VioE encapsulated to conditions with unencapsulated VioE, we calculated the concentration of VioE encapsulated in the Pdu MCPs and matched the concentration of VioE in the unencapsulated VioE condition with VioE produced in CFPS. When experiments had conditions with unencapsulated VioE and VioC, encapsulated VioE and unencapsulated VioC, and encapsulated VioE and VioC, we first measured the concentration of VioE and VioC encapsulated in Pdu MCPs. For the conditions with VioE encapsulated and VioC not encapsulated, we added Pdu MCPs with VioE encapsulated and VioC made from CFPS such that the concentrations of VioE and VioC were held constant between these two conditions. The same was done for the unencapsulated conditions where VioE and VioC made from CFPS were added to match the concentrations of VioE and VioC encapsulated in the Pdu MCP. In addition to the CFPS enzymes and the Pdu MCPs, fresh cell extract, 50 µg/mL kanamycin, 5 mM tryptophan, and 100 mM Bis Tris Buffer pH 7.2 were added to the reaction. All CFME reactions conducted in this paper were placed in 2 mL Eppendorf microcentrifuge tubes and incubated at 30 °C for 20 hours. After the reactions had been incubated, 60 µL of 25% ethyl acetate, 75% acetone was added to the reaction tube to extract the violacein products. The tube was rotated at 20 rpm at room temperature for 1 hour. The reactions were then spun down at 21,000 x g at 4°C for 10 minutes, and the supernatant was stored at -20 °C for HPLC analysis.

### HPLC

Samples were loaded onto an Agilent 1260 HPLC system in a skirted 96-well PCR plate (BioRad HSP9601). During analysis, 10 µL of sample was injected, and the metabolites were separated using a Hypersil ODS C18 HPLC column (Thermo Scientific 30105-254630) at 30 °C using 50% ethanol as the mobile phase at a flow rate of 0.2 mL/min. Metabolites were detected using a diode array detector (DAD) measuring at 574 nm. V and DV have commercially available standards, so we were able to quantify them with HPLC, whereas PV and PDV were measured in arbitrary units (AU). All of the metabolites were identified via LC-MS.

### LC-MS

Samples were injected on a 1290 Infinity II UHPLC System (Agilent Technologies Inc., Santa Clara, California, USA) onto a Poroshell 120 EC-C18 column (1.9 μm, 50 × 2.1 mm) (Agilent Technologies Inc., Santa Clara, California, USA) for C-18 chromatography which was maintained at 40 °C with a constant flow rate at 0.400 mL/min, using a gradient of mobile phase A (water, 0.1 % formic acid) and mobile phase B (100% acetonitrile, 0.1% formic acid). The gradient program was as follows: 0 – 1 min, 2 % B; 1 – 8 min, 2 – 99 % B; 8 – 10 min, hold 100 % B; 10 – 10.10 min, 99 – 2 % B; 10.10 – 15 min, hold 2 % B. “MS-Only” positive ion mode acquisition was conducted on the samples on an Agilent 6545 quadrupole time-of-flight (Q-TOF) mass spectrometer equipped with a JetStream ionization source. The source conditions were as follows: Gas Temperature, 350 °C; Drying Gas flow, 12 L/min; Nebulizer, 45 psi; Sheath Gas Temperature, 300 °C; Sheath Gas Flow, 11 L/min; VCap, 3500 V; Fragmentor, 130 V; Skimmer, 65 V; and Oct 1 RF, 750 V. The acquisition rate in MS-Only mode was 3 spectra/second, from m/z 100 – 1700 m/z range and utilizing m/z 121.05087300 and m/z 922.00979800 as reference masses which are introduced into the ion source by a separate nebulizer. The flow was maintained by an isocratic pump.

### LC-MS Data Analysis

Sample files (.d files) were imported to Agilent Mass Hunter Qualitative Analysis B10.0. The “Find by Formula” method was used to identify V, DV, PV, and PDV by matching the accurate mass of the m/z value of the [M+H] + ion and its theoretical isotopic distribution. Chromatographic peak areas were subsequently determined to calculate the relative abundance of the compounds.

## Acknowledgements

We thank and acknowledge members of the Tullman-Ercek group for their helpful discussion and preparation of this manuscript as well as constructive comments during the process of writing this manuscript. We would like to specifically acknowledge Marilyn Slininger Lee for conceiving the design for encapsulating the violacein pathway enzymes, and to Elizabeth Johnson, Madeline Joseph, Svetlana Ikonomova, Charlotte Abrahamson, and Carolyn Mills for helpful discussions.

We would also like to thank Meagan Olsen for their help with performing the autoradiograms and C^14^ leucine incorporation assays for the CFPS reactions, Fernando “Ralph” Tobias for their assistance with conducting the LC-MS to identify the violacein metabolites, and Laura Hertz for her critical figure input. We want to thank the NU IMSERC and NUANCE cores for allowing us to use their facilities for this work.

## Funding and additional information

This work was funded in part by the Army Research Office (grant W911NF-19-1-0298 to D.T.E.), the Department of Energy (grant DE-SC0022180 to D.T.E.), and National Science Foundation (grants CBET-1844336 and DMR-2308691). BJP was partially funded by a National Science Foundation Graduate training grant (grant DGE-2021900) via the Northwestern University Synthetic Biology Across the Scales Training Program.

## Conflict of interest

The authors declare that they have no conflicts of interest with the contents of this article.

## Supporting Information

## Abbreviations

1,2-PD: 1,2-propanediol
CFME: cell-free metabolic engineering
CFPS: cell-free protein synthesis
DV: Deoxyviolacein
epVioE^FL^: encapsulation peptide ssD genetically fused to the N-terminus of VioE and a FLAG epitope genetically fused to the C-terminus of VioC
epVioC^FL^: encapsulation peptide ssD genetically fused to the N-terminus of VioC and a FLAG epitope genetically fused to the C-terminus of VioC
HPLC: high-performance liquid chromatography
LC-MS: liquid chromatography with mass spectrometry
MCP: bacterial microcompartment
Pdu MCP: 1,2-propanediol bacterial microcompartment
PDV: Prodeoxyviolacein
PV: Proviolacein
SDS-PAGE: SDS-polyacrylamide gel electrophoresis
V: Violacein

---

**Figure SI1.**
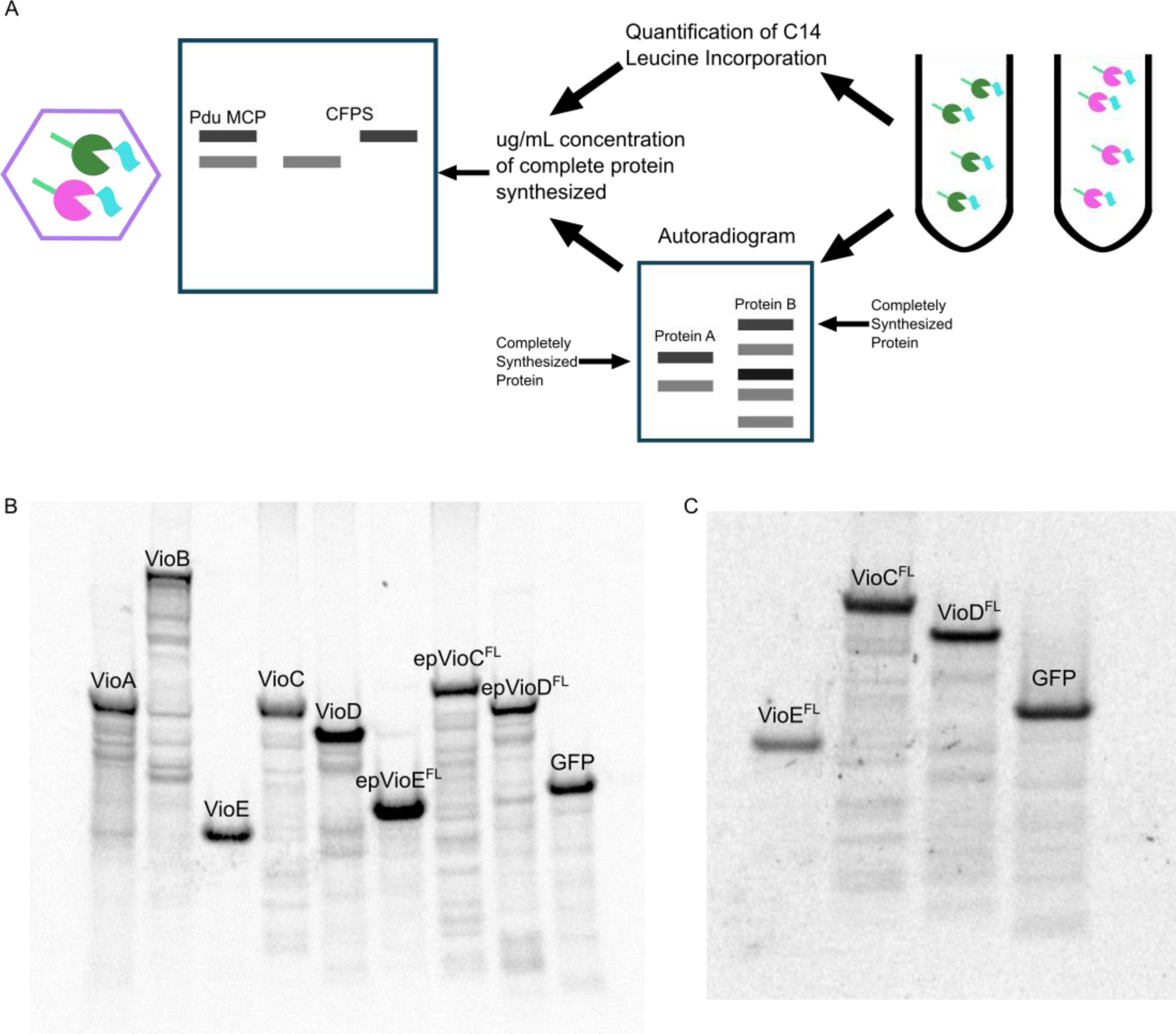
- Measuring the amount of violacein pathway enzyme encapsulated in the Pdu MCP using western blotting, SDS-PAGE autoradiograms, and ^14^Cleucine incorporation. A: A diagram of how the use of SDS-PAGE autoradiograms and ^14^C leucine incorporation were used to determine the concentration of completely synthesized violacein pathway enzyme in CFPS. The concentration of the CFPS generated enzymes was used to compare encapsulated violacein pathway enzymes to the CFPS reactions to determine the concentration of violacein pathway enzyme encapsulated in the Pdu MCP. B and C: Exposed autoradiograms of SDS-PAGE gels with CFPS reactions with radioactive ^14^C-leucine of the violacein pathway enzymes.

**Figure SI2.**
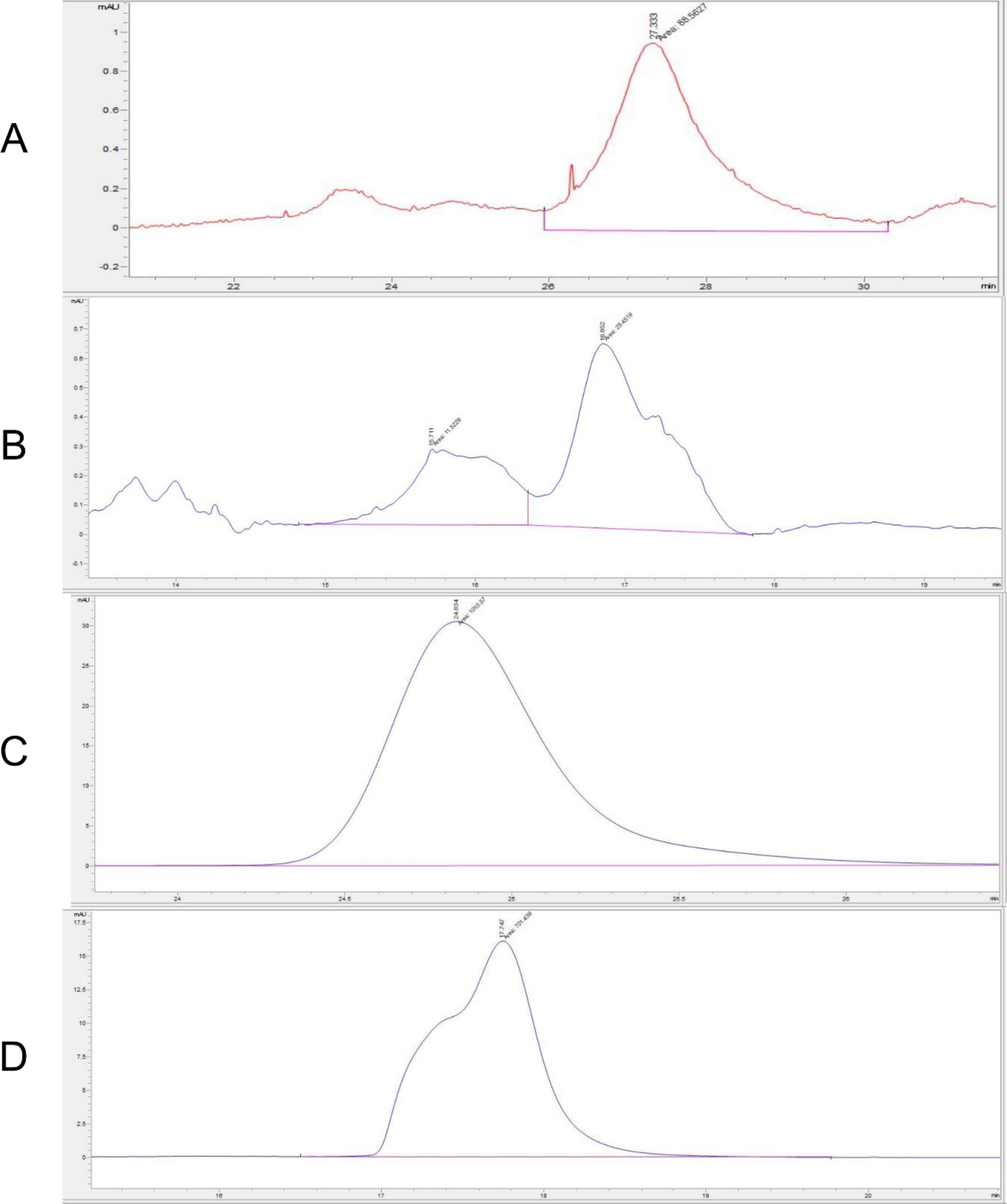
Example HPLC chromatograms for each violacein metabolite measured. Chromatograms for the violacein metabolites, PDV, PV, DV, and V were detected on an Agilent 1260 HPLC system using the Diode Array Detector at 574 nm. Retention time is on the x-axis with relative intensity on the y-axis in AU. Chromatograms taken of violacein metabolites that were produced in Figure 1C. A: Chromatogram of PDV. B: Chromatogram of PV. C: Chromatogram of DV. D: Chromatogram of V.

**Figure SI3.**
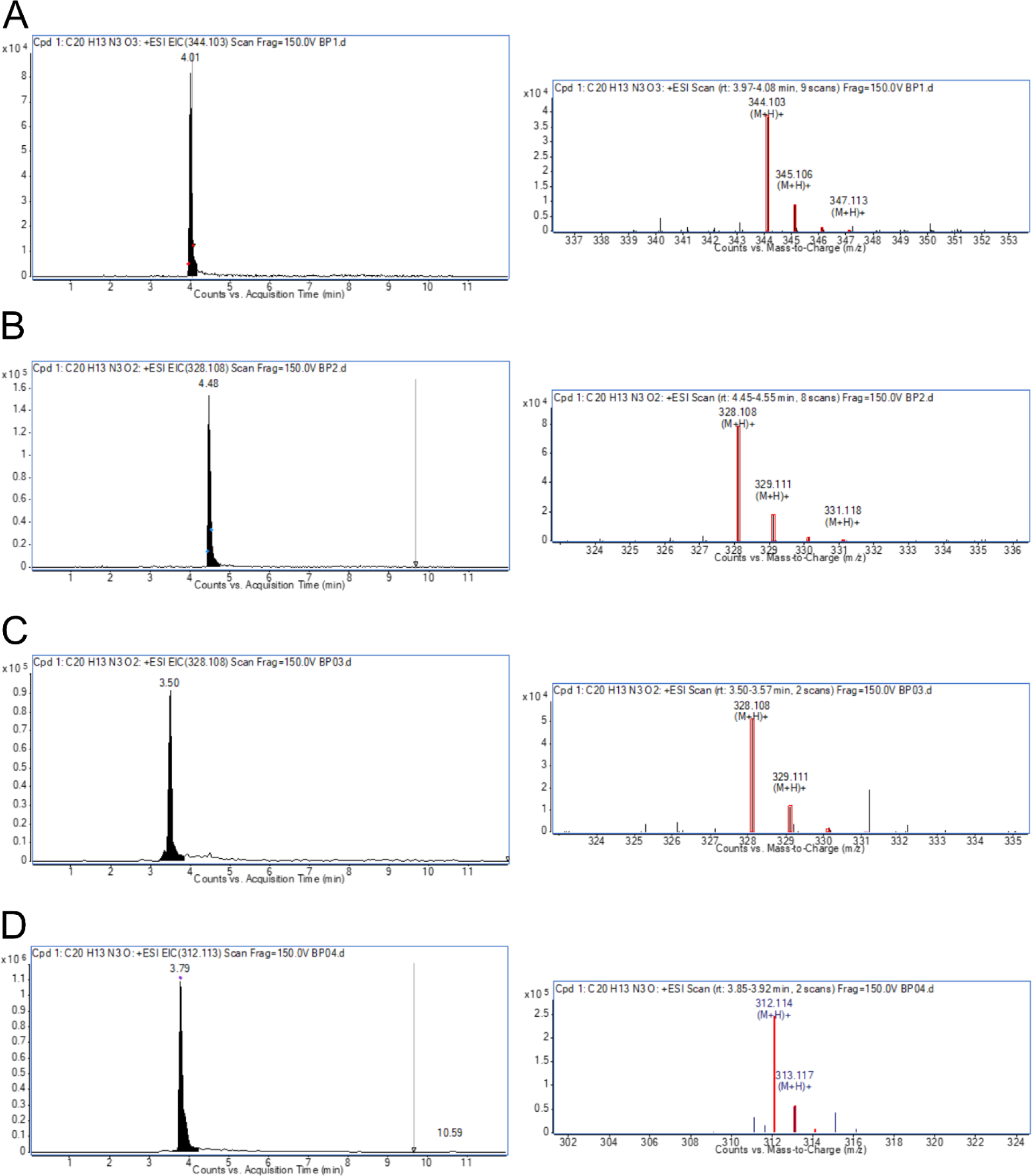
Liquid chromatography mass spectrometry confirmation of the violacein metabolites. Chromatograms and mass spectrometry for the violacein metabolites, V, DV, PV, and PDV. Chromatograms (left) with the arbitrary units on the y-axis and counts vs. acquisition time (min) on the x-axis. Mass spectrometry (right) with arbitrary units on the y-axis and counts vs. mass-to-charge (m/z) on the x-axis. Red boxes indicate the theoretical isotope pattern of the chemical formula of the compound. A: Violacein. B: Deoxyviolacein. C: Proviolacein. D: Prodeoxyviolacein.

**Figure SI4.**
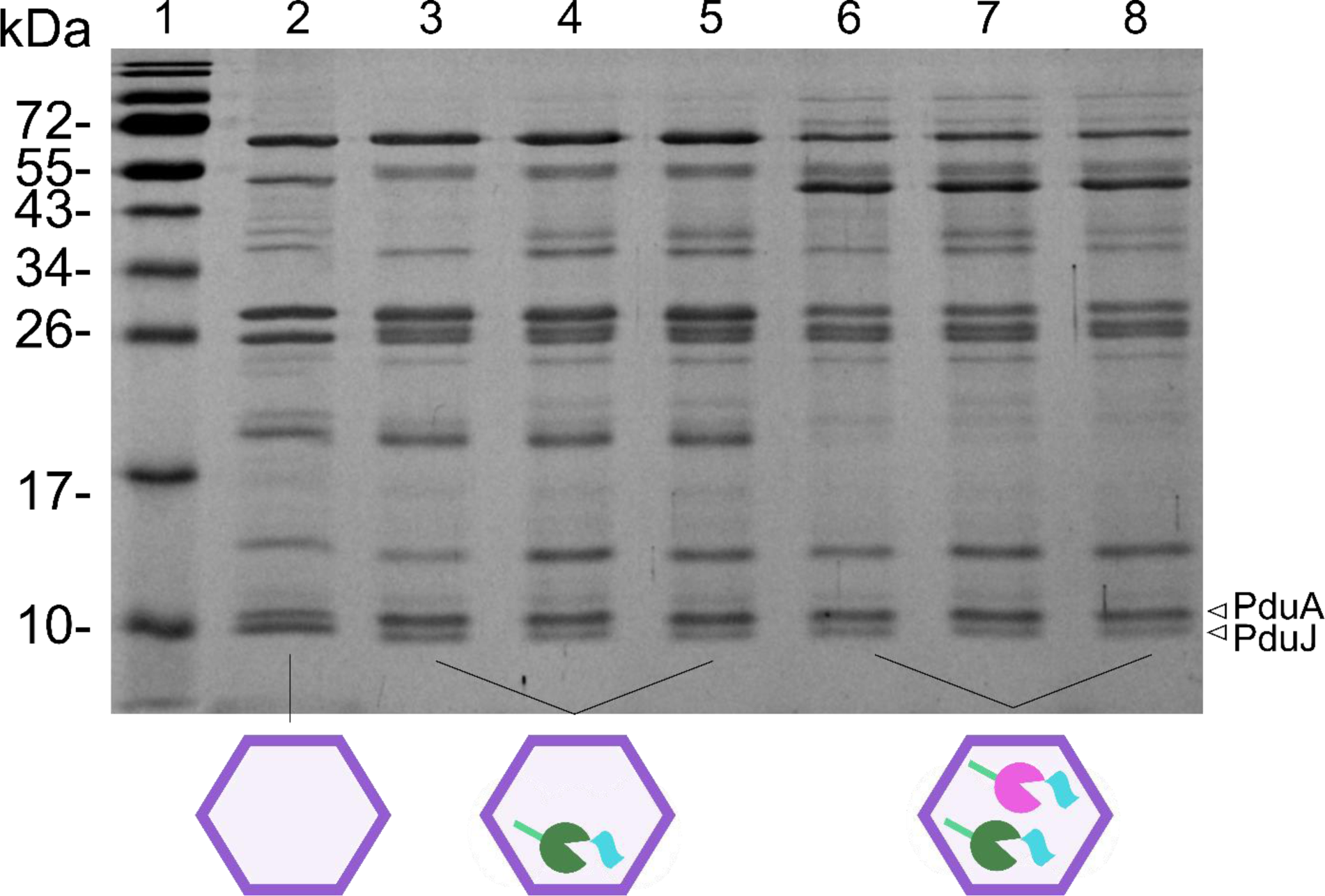
Coomassie Blue stain of SDS-PAGE gels with the purified Pdu MCPs. Purified Pdu MCPs run on an SDS-PAGE gel that was then stained with Coomassie Blue stain. Lane 1: Fisher BioReagents EZ-Run Prestained Rec Protein Ladder, Lane 2: WT Pdu MCPs, Lanes 3-5: Biological replicates (N = 3) of Pdu MCPs with epVioE^FL^ encapsulated, Lanes 6-8: Biological replicates (N = 3) of Pdu MCPs with epVioC^FL^ encapsulated. At around 10 kDa, bands at the expected size for PduA and PduJ were observed, suggesting that these are indeed purified Pdu MCPs.

**Figure SI5.**
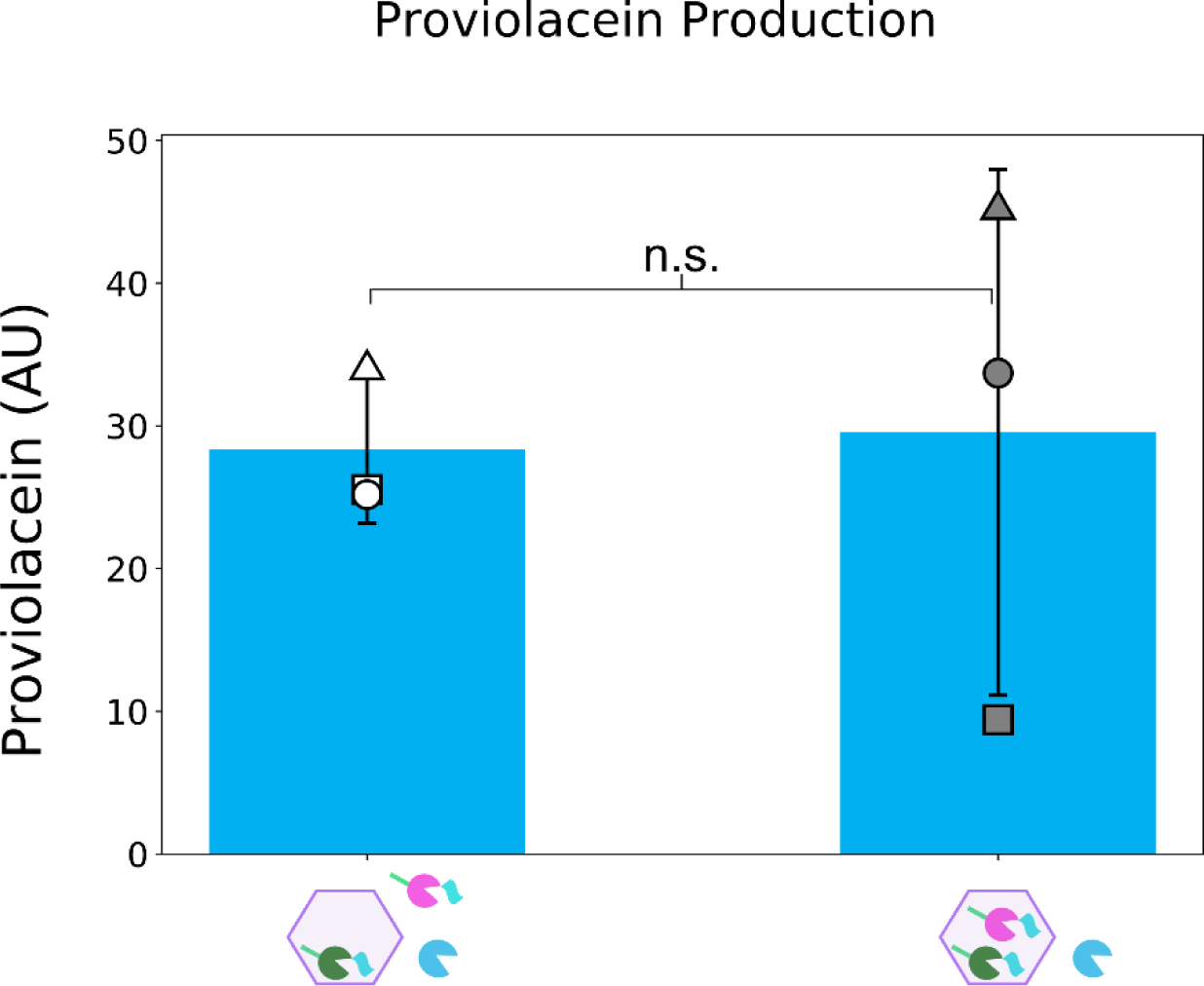
Production of violacein pathway product proviolacein in CFME reactions with Pdu MCPs with epVioE^FL^ and epVioC^FL^ encapsulated. Production of PV in CFME reactions with VioD not encapsulated and with Pdu MCPs with epVioE^FL^ encapsulated and epVioC^FL^ not encapsulated or with Pdu MCPs with epVioE^FL^ and epVioC^FL^ encapsulated. All CFME reactions have VioA and VioB added but not depicted. Error bars denote +/- one standard deviation from biological replicates (N = 3). The encapsulated epVioE^FL^ and not encapsulated epVioC^FL^ condition is denoted with white-filled symbols, and the encapsulated epVioC^FL^ and epVioC^FL^ condition is denoted with grey filled symbols. The first replicate is represented by a □, the second replicate is represented by a Δ, and the third replicate is represented by a O. Significance was determined with a two tailed *t*-test and is denoted with the following: * - p < 0.01, ** - p < 0.05, *** - p < 0.01, n.s. – p > 0.1.

**Table SI1.**
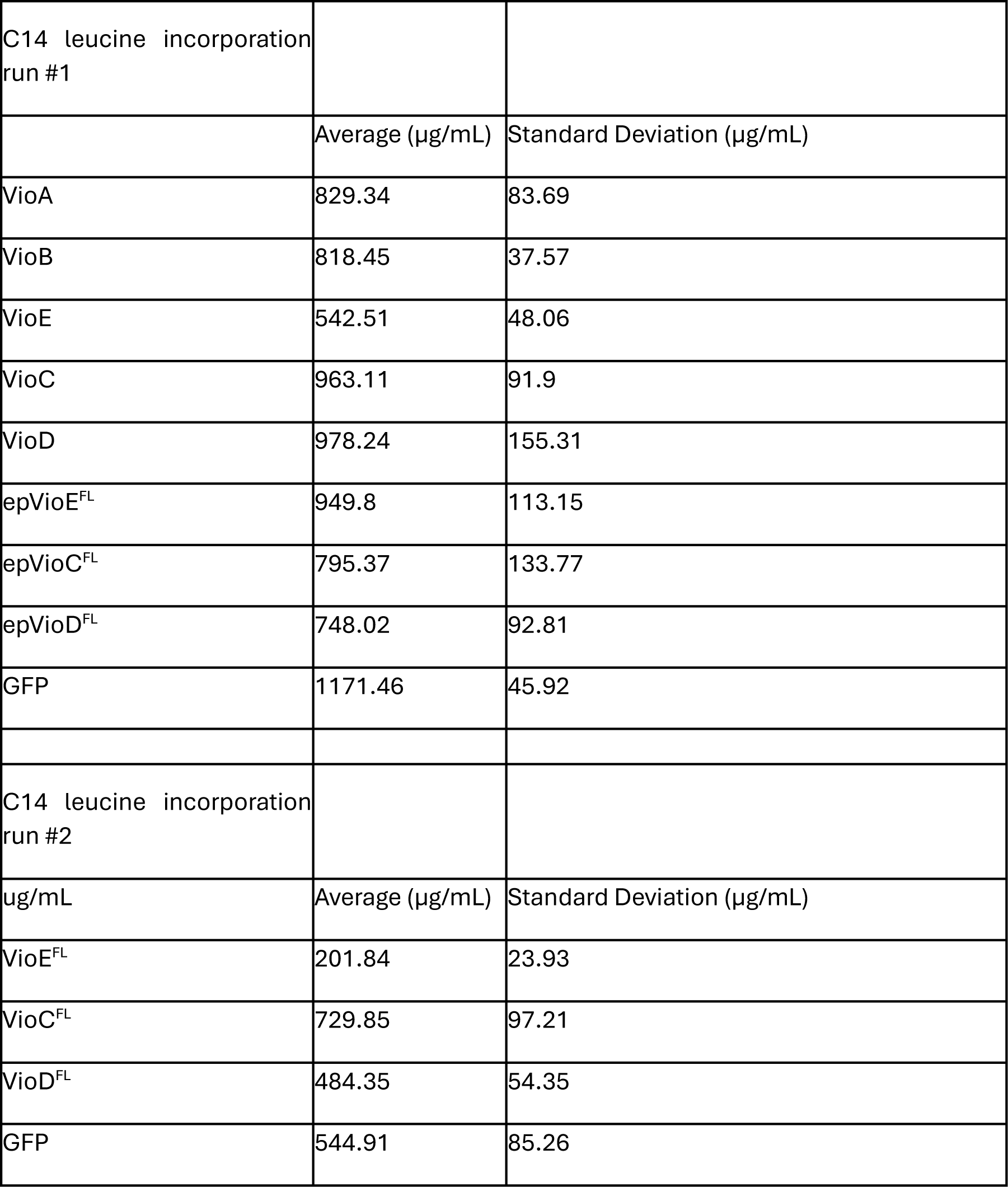
- Protein production in CFPS: Protein concentrations of the violacein pathway enzymes made in CFPS reactions with ^14^C-Leucine added of technical replicates (N = 3).

|  |  |  |
| --- | --- | --- |
| C14 leucine incorporation<br>run #1 |  |  |
|  | Average (µg/mL) | Standard Deviation (µg/mL) |
| VioA | 829.34 | 83.69 |
| VioB | 818.45 | 37.57 |
| VioE | 542.51 | 48.06 |
| VioC | 963.11 | 91.9 |
| VioD | 978.24 | 155.31 |
| epVioE<sup>FL</sup> | 949.8 | 113.15 |
| epVioC<sup>FL</sup> | 795.37 | 133.77 |
| epVioD<sup>FL</sup> | 748.02 | 92.81 |
| GFP | 1171.46 | 45.92 |
| C14 leucine incorporation<br>run #2 |  |  |
| ug/mL | Average (µg/mL) | Standard Deviation (µg/mL) |
| VioE<sup>FL</sup> | 201.84 | 23.93 |
| VioC<sup>FL</sup> | 729.85 | 97.21 |
| VioD<sup>FL</sup> | 484.35 | 54.35 |
| GFP | 544.91 | 85.26 |

**Table SI2.**
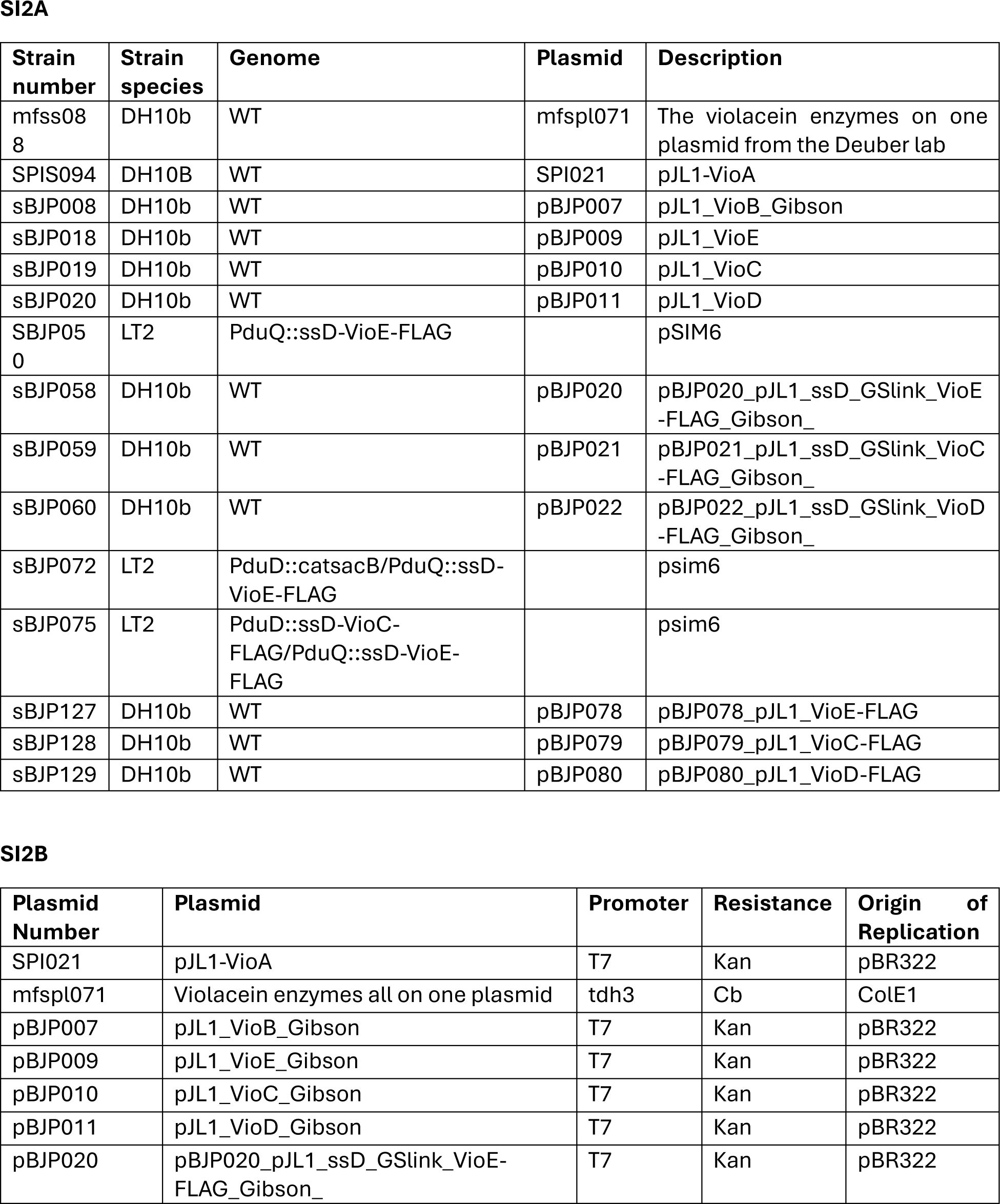
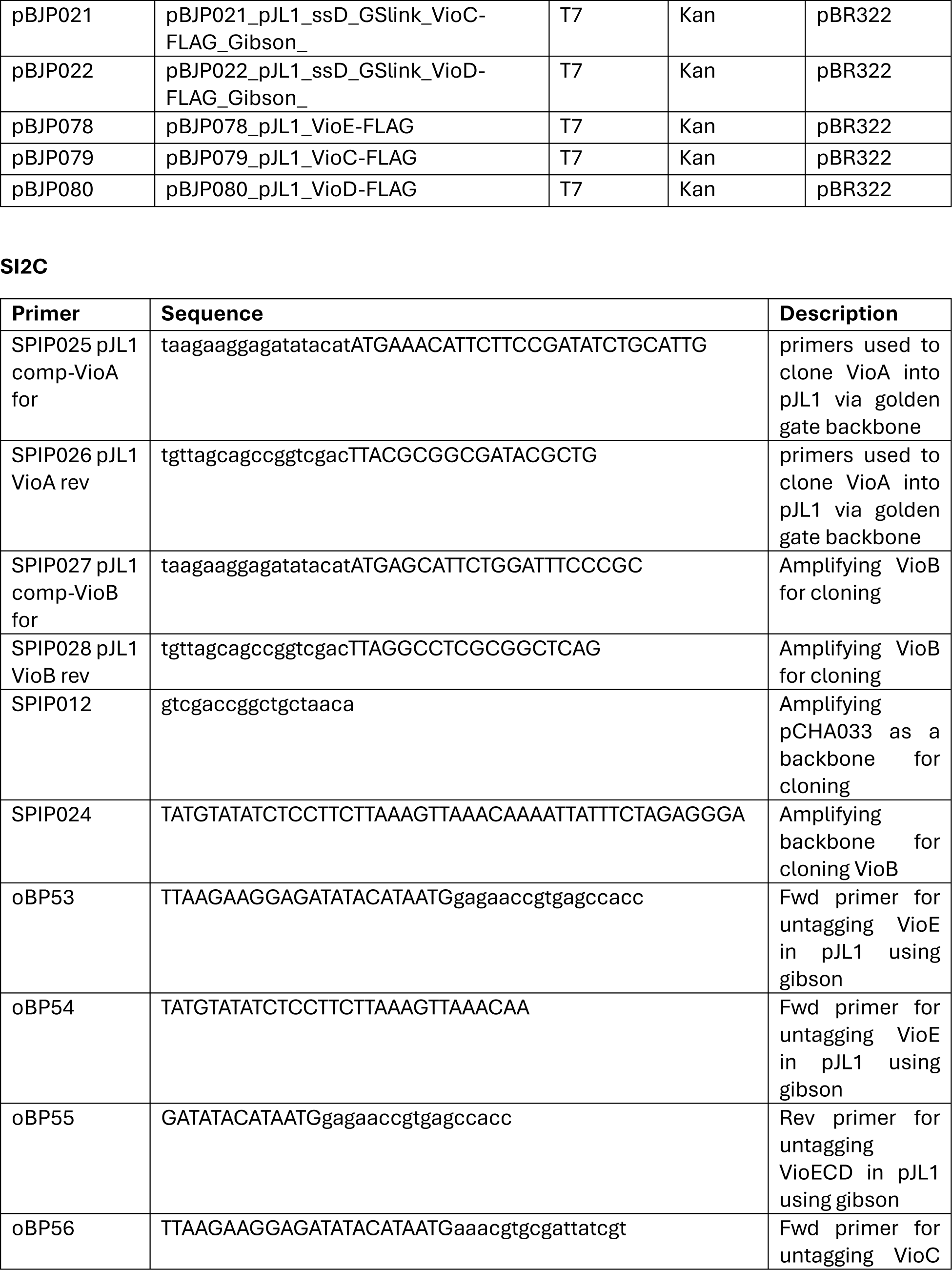
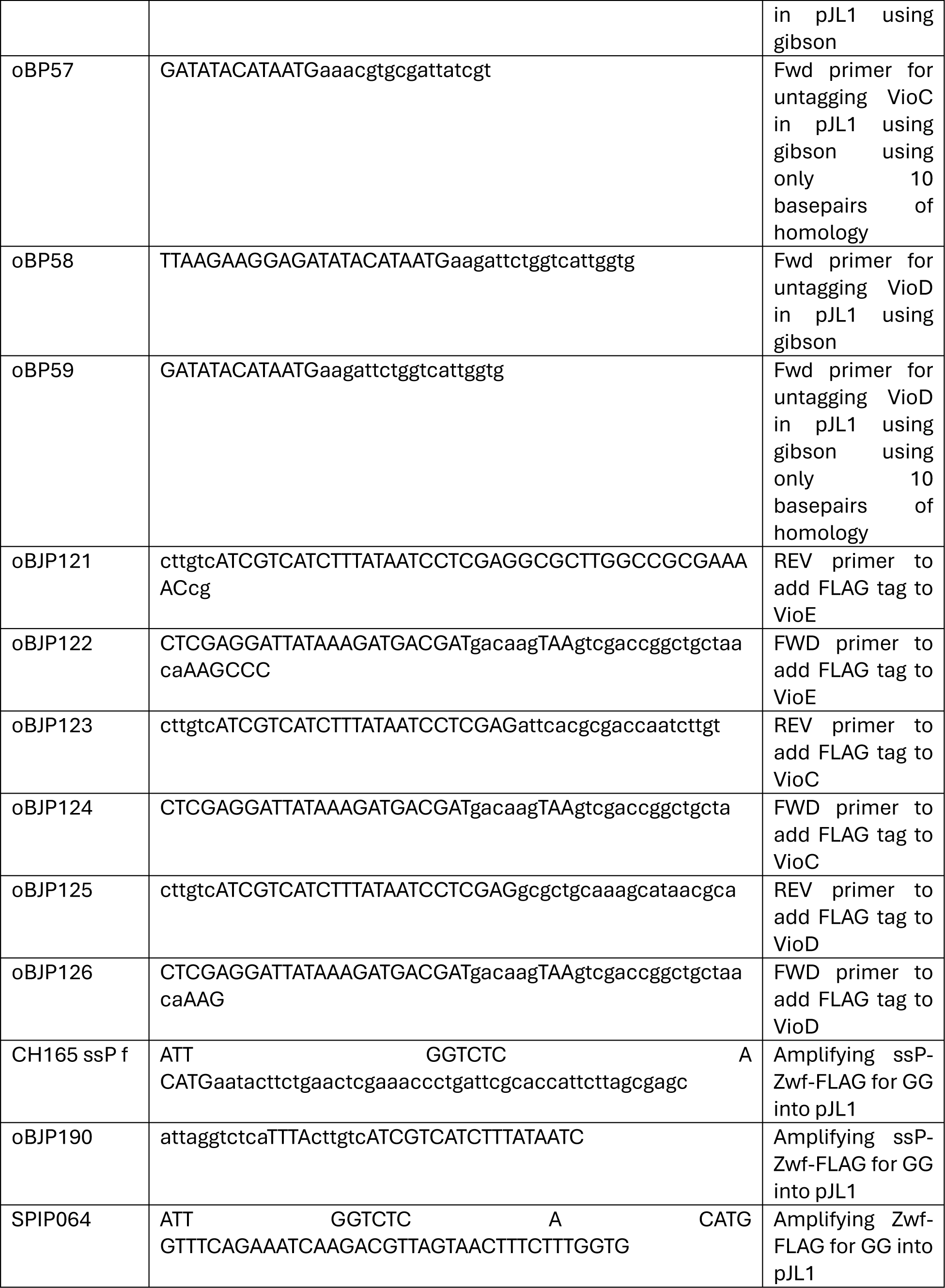
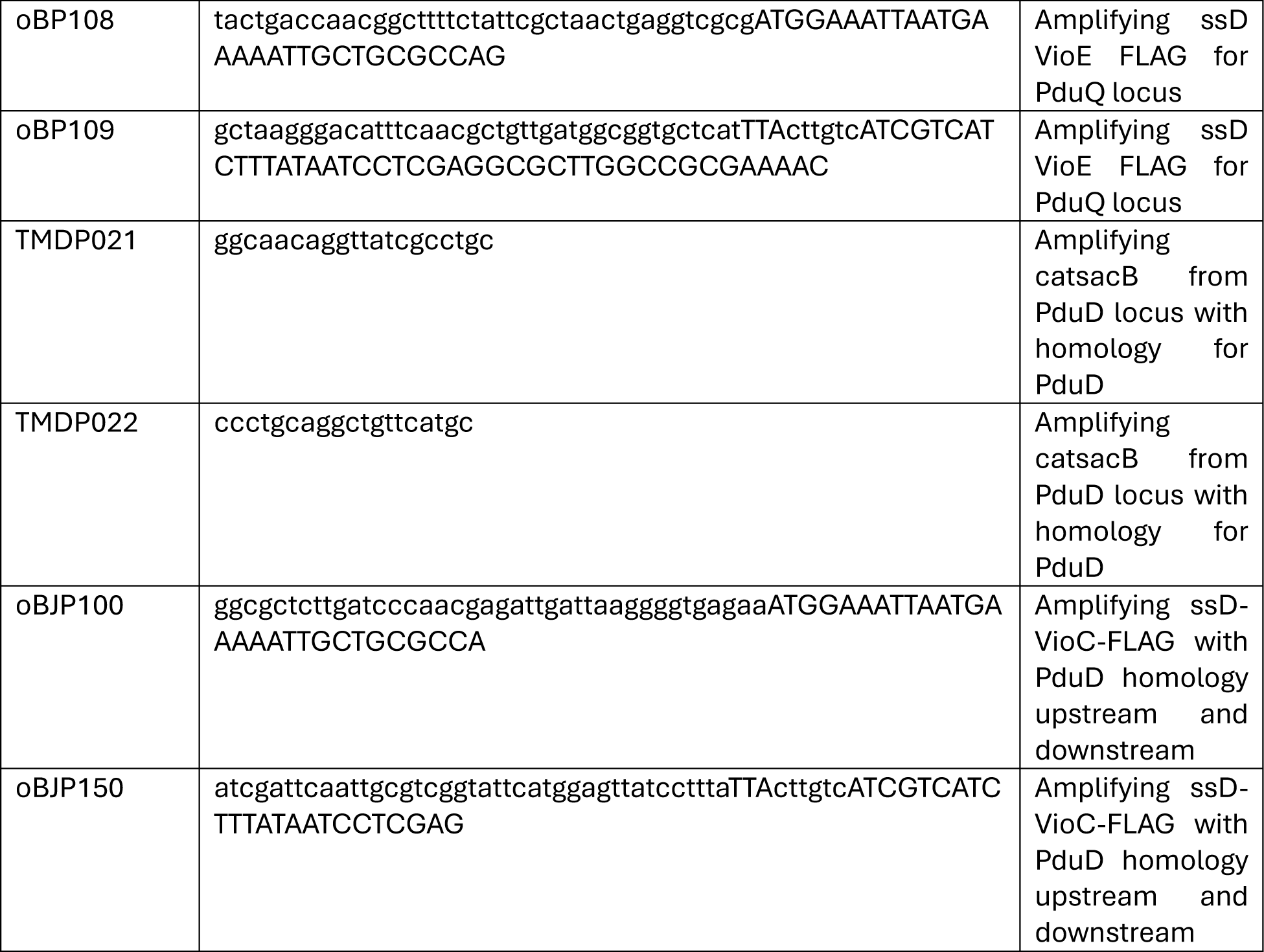
Strains, plasmids, and primers used in this work: A: Table of the strains utilized in this work for cloning and Pdu MCP purification. B: Plasmids utilized in this work for protein production in CFPS. C: Primers utilized in this work for cloning and recombineering.

## Notes

### Competing Interest Statement

The authors have declared no competing interest.

