## Supplemental Information for "Encapsulation of Select Violacein Pathway Enzymes in the 1,2-Propanediol Utilization Bacterial Microcompartment to Divert Pathway Flux"

**Supporting Information**

**Table SI1 - Protein production in CFPS:** Protein concentrations of the violacein pathway enzymes made in CFPS reactions with $^{14}$C-Leucine added of technical replicates (N = 3).

| | | |
|---|---|---|
| C14 leucine incorporation run #1 | | |
| | Average (μg/mL) | Standard Deviation (μg/mL) |
| VioA | 829.34 | 83.69 |
| VioB | 818.45 | 37.57 |
| VioE | 542.51 | 48.06 |
| VioC | 963.11 | 91.9 |
| VioD | 978.24 | 155.31 |
| epVioE$^{FL}$ | 949.8 | 113.15 |
| epVioC$^{FL}$ | 795.37 | 133.77 |
| epVioD$^{FL}$ | 748.02 | 92.81 |
| GFP | 1171.46 | 45.92 |
| C14 leucine incorporation run #2 | | |
| ug/mL | Average (μg/mL) | Standard Deviation (μg/mL) |
| VioE$^{FL}$ | 201.84 | 23.93 |
| VioC$^{FL}$ | 729.85 | 97.21 |
| VioD$^{FL}$ | 484.35 | 54.35 |
| GFP | 544.91 | 85.26 |

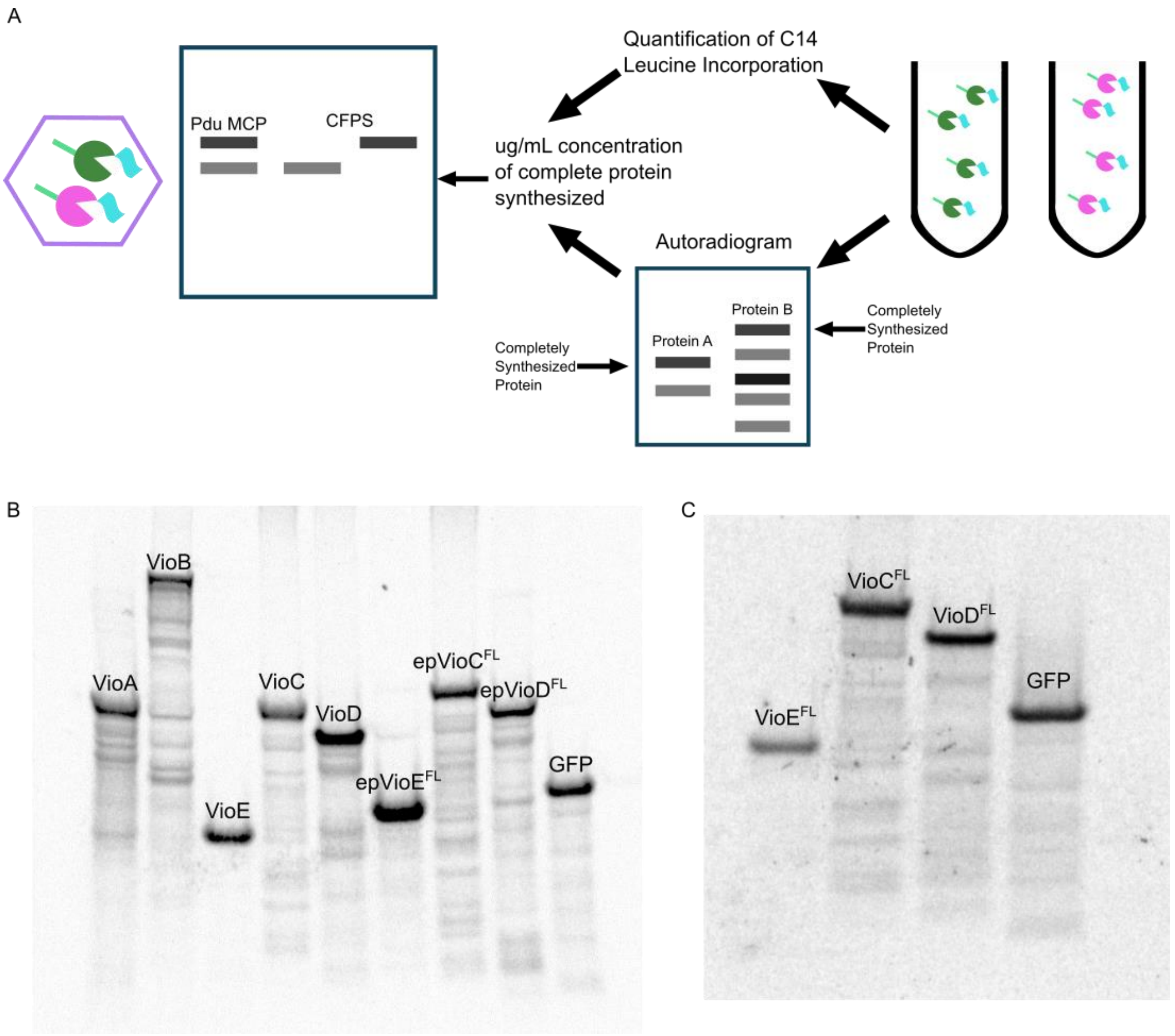

**Figure SI1 - Measuring the amount of violacein pathway enzyme encapsulated in the Pdu MCP using western blotting, SDS-PAGE autoradiograms, and $^{14}$Cleucine incorporation.** A: A diagram of how the use of SDS-PAGE autoradiograms and $^{14}$C leucine incorporation were used to determine the concentration of completely synthesized violacein pathway enzyme in CFPS. The concentration of the CFPS generated enzymes was used to compare encapsulated violacein pathway enzymes to the CFPS reactions to determine the concentration of violacein pathway enzyme encapsulated in the Pdu MCP. B and C: Exposed autoradiograms of SDS-PAGE gels with CFPS reactions with radioactive $^{14}$C-leucine of the violacein pathway enzymes.

**Table SI2 – Strains, plasmids, and primers used in this work:** A: Table of the strains utilized in this work for cloning and Pdu MCP purification. B: Plasmids utilized in this work for protein production in CFPS. C: Primers utilized in this work for cloning and recombineering.

**SI2A**

| Strain number | Strain species | Genome | Plasmid | Description |
|---|---|---|---|---|
| mfss08 8 | DH10b | WT | mfspl071 | The violacein enzymes on one plasmid from the Deuber lab |
| SPIS094 | DH10B | WT | SPI021 | pJL1-VioA |
| sBJP008 | DH10b | WT | pBJP007 | pJL1_VioB_Gibson |
| sBJP018 | DH10b | WT | pBJP009 | pJL1_VioE |
| sBJP019 | DH10b | WT | pBJP010 | pJL1_VioC |
| sBJP020 | DH10b | WT | pBJP011 | pJL1_VioD |
| SBJP05 0 | LT2 | PduQ::ssD-VioE-FLAG | | pSIM6 |
| sBJP058 | DH10b | WT | pBJP020 | pBJP020_pJL1_ssD_GSlink_VioE -FLAG_Gibson_ |
| sBJP059 | DH10b | WT | pBJP021 | pBJP021_pJL1_ssD_GSlink_VioC -FLAG_Gibson_ |
| sBJP060 | DH10b | WT | pBJP022 | pBJP022_pJL1_ssD_GSlink_VioD -FLAG_Gibson_ |
| sBJP072 | LT2 | PduD::catsacB/PduQ::ssD-VioE-FLAG | | psim6 |
| sBJP075 | LT2 | PduD::ssD-VioC-FLAG/PduQ::ssD-VioE-FLAG | | psim6 |
| sBJP127 | DH10b | WT | pBJP078 | pBJP078_pJL1_VioE-FLAG |
| sBJP128 | DH10b | WT | pBJP079 | pBJP079_pJL1_VioC-FLAG |
| sBJP129 | DH10b | WT | pBJP080 | pBJP080_pJL1_VioD-FLAG |

**SI2B**

| Plasmid Number | Plasmid | Promoter | Resistance | Origin of Replication |
|---|---|---|---|---|
| SPI021 | pJL1-VioA | T7 | Kan | pBR322 |
| mfspl071 | Violacein enzymes all on one plasmid | tdh3 | Cb | ColE1 |
| pBJP007 | pJL1_VioB_Gibson | T7 | Kan | pBR322 |
| pBJP009 | pJL1_VioE_Gibson | T7 | Kan | pBR322 |
| pBJP010 | pJL1_VioC_Gibson | T7 | Kan | pBR322 |
| pBJP011 | pJL1_VioD_Gibson | T7 | Kan | pBR322 |
| pBJP020 | pBJP020_pJL1_ssD_GSlink_VioE-FLAG_Gibson_ | T7 | Kan | pBR322 |

| | | | | |
|---|---|---|---|---|
| pBJP021 | pBJP021_pJL1_ssD_GSlink_VioC-FLAG_Gibson_ | T7 | Kan | pBR322 |
| pBJP022 | pBJP022_pJL1_ssD_GSlink_VioD-FLAG_Gibson_ | T7 | Kan | pBR322 |
| pBJP078 | pBJP078_pJL1_VioE-FLAG | T7 | Kan | pBR322 |
| pBJP079 | pBJP079_pJL1_VioC-FLAG | T7 | Kan | pBR322 |
| pBJP080 | pBJP080_pJL1_VioD-FLAG | T7 | Kan | pBR322 |

**SI2C**

| **Primer** | **Sequence** | **Description** |
|---|---|---|
| SPIP025 pJL1 comp-VioA for | taagaaggagatatacatATGAAACATTCTTCCGATATCTGCATTG | primers used to clone VioA into pJL1 via golden gate backbone |
| SPIP026 pJL1 VioA rev | tgttagcagccggtcgacTTACGCGGCGATACGCTG | primers used to clone VioA into pJL1 via golden gate backbone |
| SPIP027 pJL1 comp-VioB for | taagaaggagatatacatATGAGCATTCTGGATTTCCCGC | Amplifying VioB for cloning |
| SPIP028 pJL1 VioB rev | tgttagcagccggtcgacTTAGGCCTCGCGGCTCAG | Amplifying VioB for cloning |
| SPIP012 | gtcgaccggctgctaaca | Amplifying pCHA033 as a backbone for cloning |
| SPIP024 | TATGTATATCTCCTTCTTAAAGTTAAACAAAATTATTTCTAGAGGGA | Amplifying backbone for cloning VioB |
| oBP53 | TTAAGAAGGAGATATACATAATGgagaaccgtgagccacc | Fwd primer for untagging VioE in pJL1 using gibson |
| oBP54 | TATGTATATCTCCTTCTTAAAGTTAAACAA | Fwd primer for untagging VioE in pJL1 using gibson |
| oBP55 | GATATACATAATGgagaaccgtgagccacc | Rev primer for untagging VioECD in pJL1 using gibson |
| oBP56 | TTAAGAAGGAGATATACATAATGaaacgtgcgattatcgt | Fwd primer for untagging VioC |

| | | |
|---|---|---|
| | | in pJL1 using gibson |
| oBP57 | GATATACATAATGaaacgtgcgattatcgt | Fwd primer for untagging VioC in pJL1 using gibson using only 10 basepairs of homology |
| oBP58 | TTAAGAAGGAGATATACATAATGaagattctggtcattggtg | Fwd primer for untagging VioD in pJL1 using gibson |
| oBP59 | GATATACATAATGaagattctggtcattggtg | Fwd primer for untagging VioD in pJL1 using gibson using only 10 basepairs of homology |
| oBJP121 | cttgtcATCGTCATCTTTATAATCCTCGAGGCGCTTGGCCGCGAAAACcg | REV primer to add FLAG tag to VioE |
| oBJP122 | CTCGAGGATTATAAAGATGACGATgacaagTAAgtcgaccggctgctaacaAAGCCC | FWD primer to add FLAG tag to VioE |
| oBJP123 | cttgtcATCGTCATCTTTATAATCCTCGAGattcacgcgaccaatcttgt | REV primer to add FLAG tag to VioC |
| oBJP124 | CTCGAGGATTATAAAGATGACGATgacaagTAAgtcgaccggctgcta | FWD primer to add FLAG tag to VioC |
| oBJP125 | cttgtcATCGTCATCTTTATAATCCTCGAGgcgctgcaaagcataacgca | REV primer to add FLAG tag to VioD |
| oBJP126 | CTCGAGGATTATAAAGATGACGATgacaagTAAgtcgaccggctgctaacaAAG | FWD primer to add FLAG tag to VioD |
| CH165 ssP f | ATT GGTCTC A CATGaatacttctgaactcgaaaccctgattcgcaccattcttagcgagc | Amplifying ssP-Zwf-FLAG for GG into pJL1 |
| oBJP190 | attaggtctcaTTTActtgtcATCGTCATCTTTATAATC | Amplifying ssP-Zwf-FLAG for GG into pJL1 |
| SPIP064 | ATT GGTCTC A CATG GTTTCAGAAATCAAGACGTTAGTAACTTTCTTTGGTG | Amplifying Zwf-FLAG for GG into pJL1 |

| | | |
|---|---|---|
| oBP108 | tactgaccaacggcttttctattcgctaactgaggtcgcgATGGAAATTAATGAAAAATTGCTGCGCCAG | Amplifying ssD VioE FLAG for PduQ locus |
| oBP109 | gctaagggacatttcaacgctgttgatggcggtgctcatTTActtgtcATCGTCATCTTTATAATCCTCGAGGCGCTTGGCCGCGAAAAC | Amplifying ssD VioE FLAG for PduQ locus |
| TMDP021 | ggcaacaggttatcgcctgc | Amplifying catsacB from PduD locus with homology for PduD |
| TMDP022 | ccctgcaggctgttcatgc | Amplifying catsacB from PduD locus with homology for PduD |
| oBJP100 | ggcgctcttgatcccaacgagattgattaaggggtgagaaATGGAAATTAATGAAAAATTGCTGCGCCA | Amplifying ssD-VioC-FLAG with PduD homology upstream and downstream |
| oBJP150 | atcgattcaattgcgtcggtattcatggagttatcctttaTTActtgtcATCGTCATCTTTATAATCCTCGAG | Amplifying ssD-VioC-FLAG with PduD homology upstream and downstream |

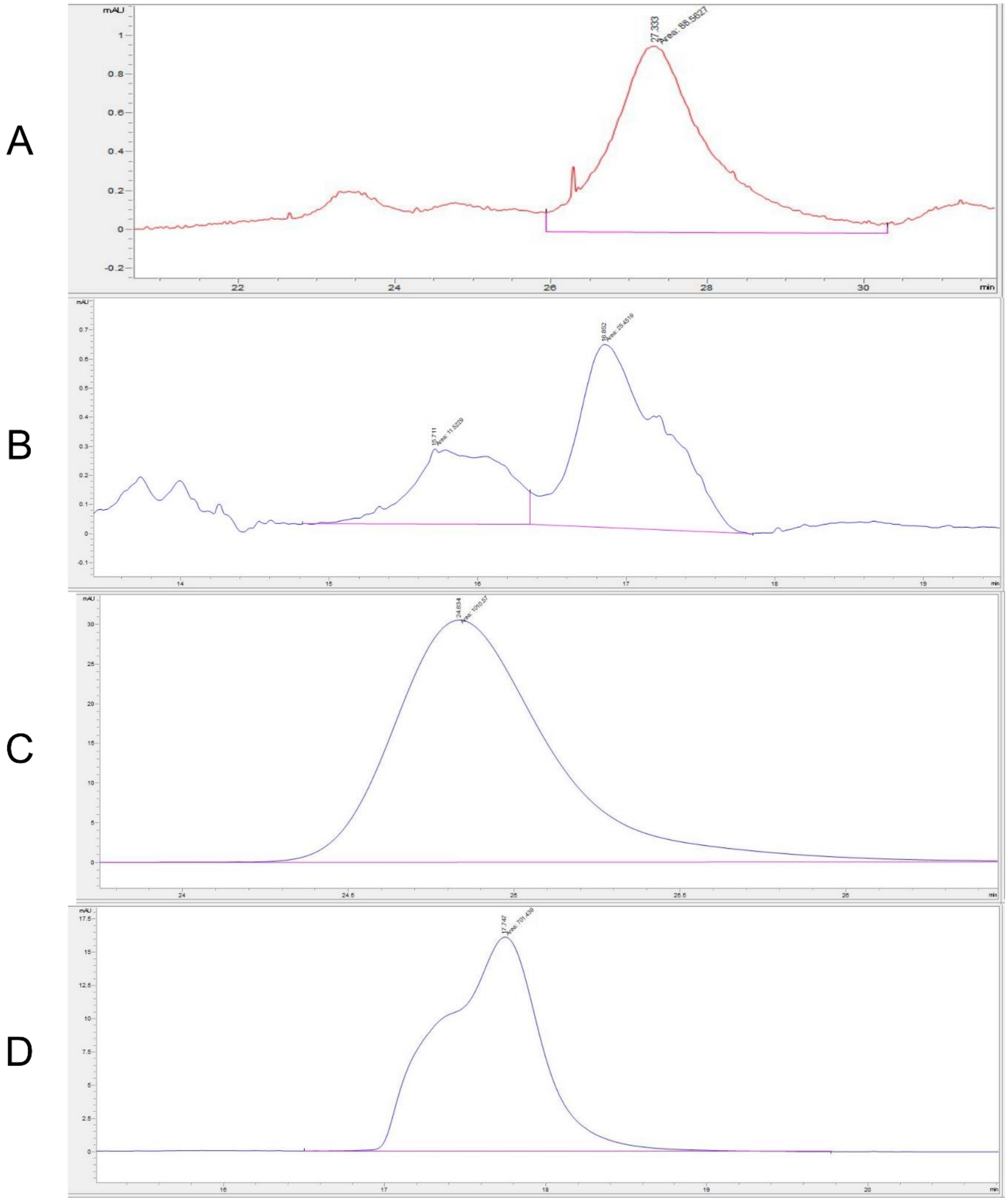

**Figure SI2 – Example HPLC chromatograms for each violacein metabolite measured.** Chromatograms for the violacein metabolites, PDV, PV, DV, and V were detected on an Agilent 1260 HPLC system using the Diode Array Detector at 574 nm. Retention time is on the x-axis with relative intensity on the y-axis in AU. Chromatograms taken of violacein metabolites that were produced in Figure 1C. A: Chromatogram of PDV. B: Chromatogram of PV. C: Chromatogram of DV. D: Chromatogram of V.

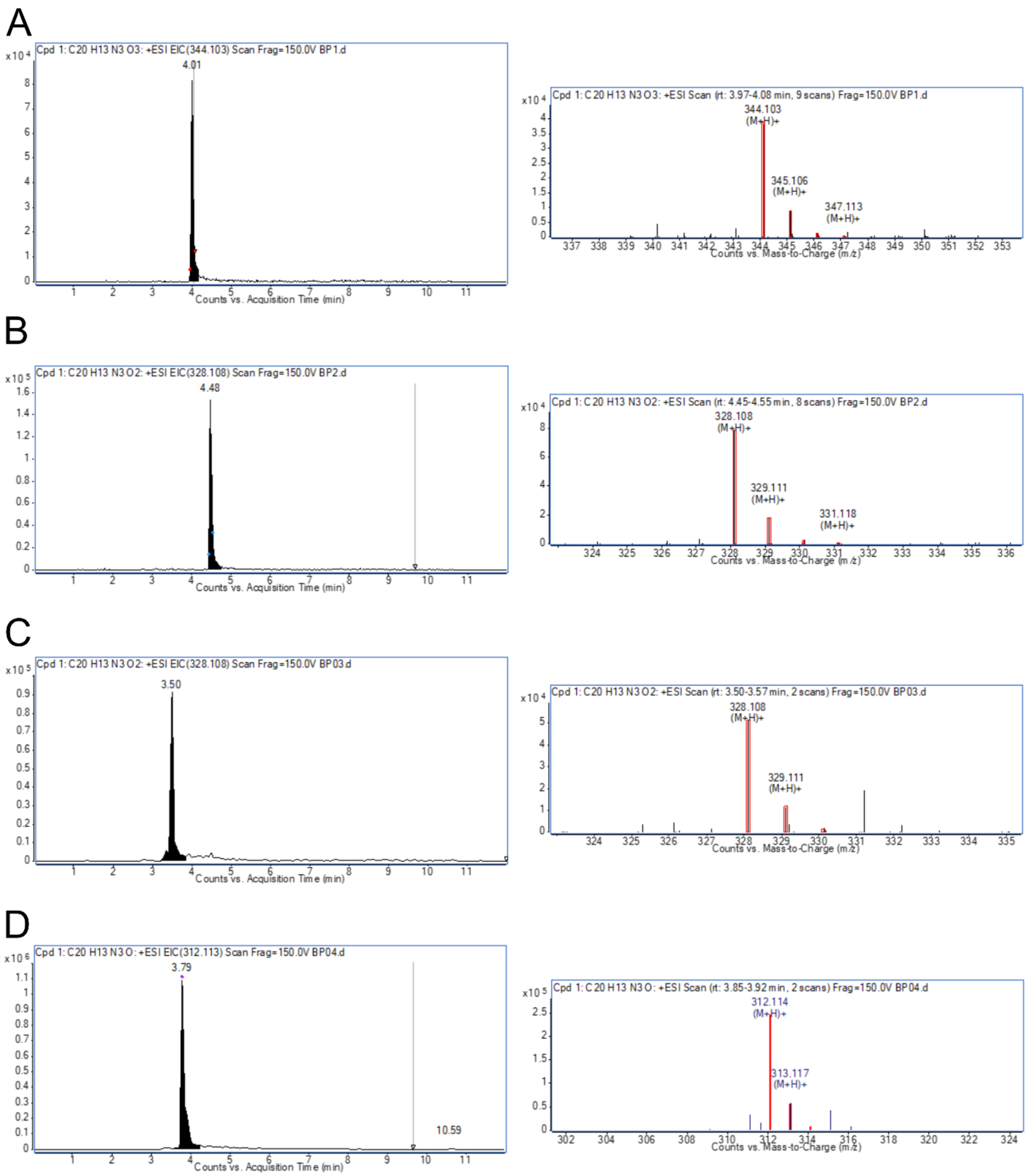

**Figure SI3 – Liquid chromatography mass spectrometry confirmation of the violacein metabolites.** Chromatograms and mass spectrometry for the violacein metabolites, V, DV, PV, and PDV. Chromatograms (left) with the arbitrary units on the y-axis and counts vs. acquisition time (min) on the x-axis. Mass spectrometry (right) with arbitrary units on the y-axis and counts vs. mass-to-charge (m/z) on the x-axis. Red boxes indicate the theoretical isotope pattern of the chemical formula of the compound. A: Violacein. B: Deoxyviolacein. C: Proviolacein. D: Prodeoxyviolacein.

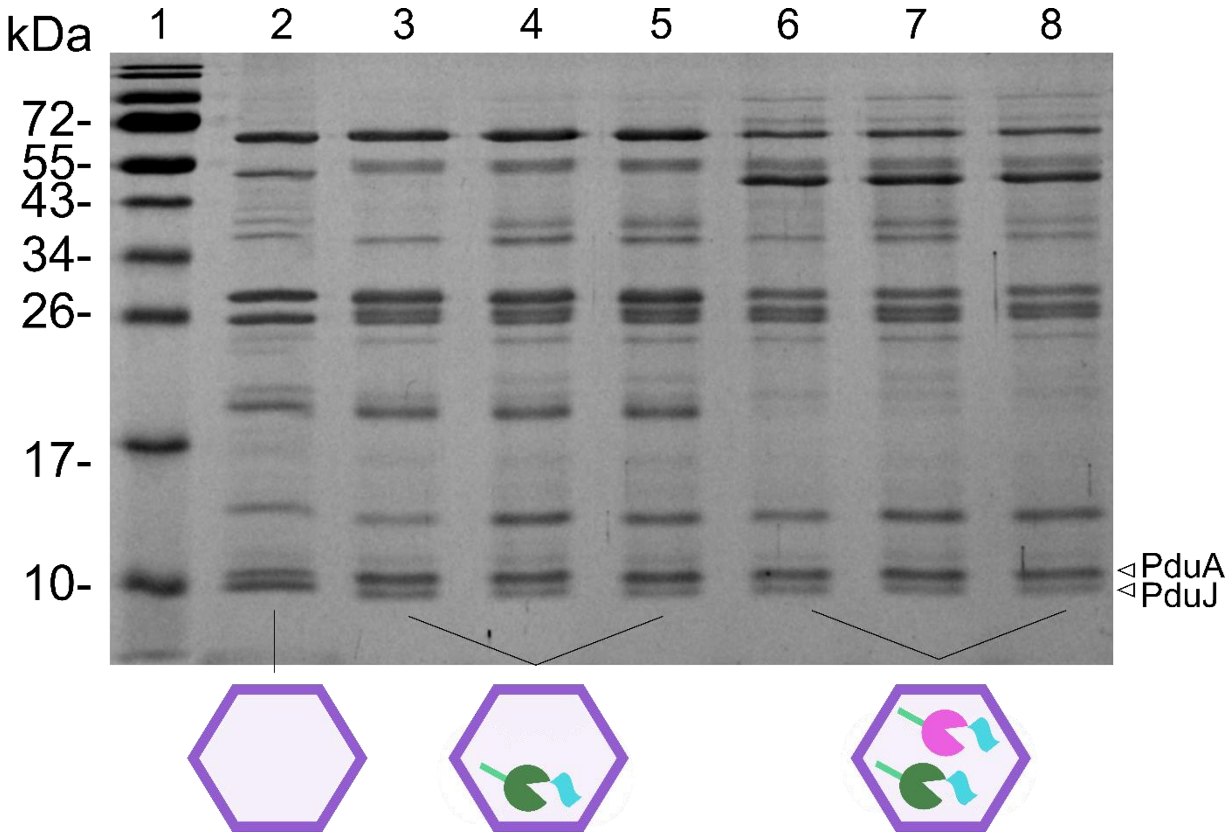

**Figure SI4 – Coomassie Blue stain of SDS-PAGE gels with the purified Pdu MCPs.** Purified Pdu MCPs run on an SDS-PAGE gel that was then stained with Coomassie Blue stain. Lane 1: Fisher BioReagents EZ-Run Prestained Rec Protein Ladder, Lane 2: WT Pdu MCPs, Lanes 3-5: Biological replicates (N = 3) of Pdu MCPs with epVioE$^{FL}$ encapsulated, Lanes 6-8: Biological replicates (N = 3) of Pdu MCPs with epVioC$^{FL}$ encapsulated. At around 10 kDa, bands at the expected size for PduA and PduJ were observed, suggesting that these are indeed purified Pdu MCPs.

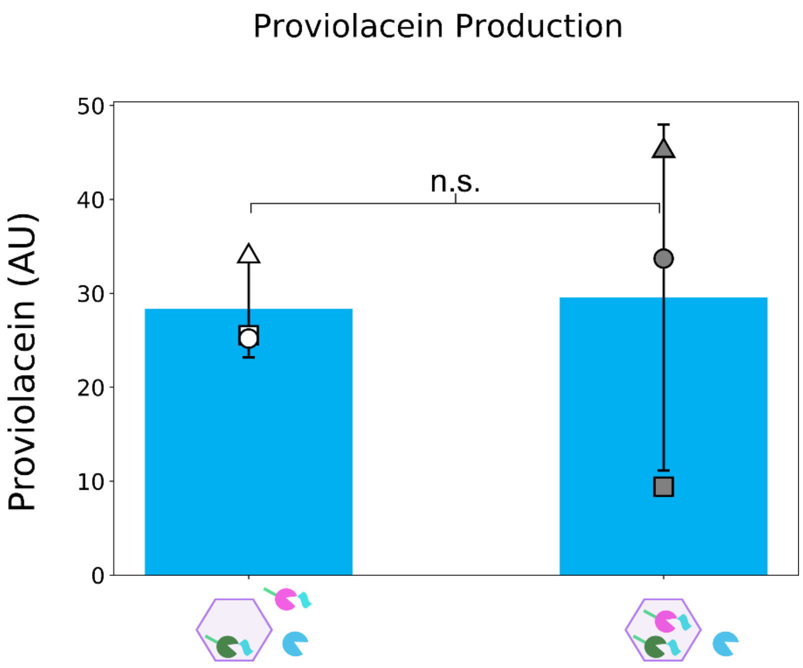

**Figure SI5 – Production of violacein pathway product proviolacein in CFME reactions with Pdu MCPs with epVioE$^{FL}$ and epVioC$^{FL}$ encapsulated.** Production of PV in CFME reactions with VioD not encapsulated and with Pdu MCPs with epVioE$^{FL}$ encapsulated and epVioC$^{FL}$ not encapsulated or with Pdu MCPs with epVioE$^{FL}$ and epVioC$^{FL}$ encapsulated. All CFME reactions have VioA and VioB added but not depicted. Error bars denote +/- one standard deviation from biological replicates (N = 3). The encapsulated epVioE$^{FL}$ and not encapsulated epVioC$^{FL}$ condition is denoted with white-filled symbols, and the encapsulated epVioC$^{FL}$ and epVioC$^{FL}$ condition is denoted with grey filled symbols. The first replicate is represented by a □, the second replicate is represented by a Δ, and the third replicate is represented by a O. Significance was determined with a two tailed *t*-test and is denoted with the following: * - $p < 0.01$, ** - $p < 0.05$, *** - $p < 0.01$, n.s. – $p > 0.1$.
